# Drosulfakinin signaling modulates female sexual receptivity in *Drosophila*

**DOI:** 10.1101/2021.12.09.471924

**Authors:** Tao Wang, Biyang Jing, Bowen Deng, Kai Shi, Jing Li, Baoxu Ma, Fengming Wu, Chuan Zhou

## Abstract

Female sexual behavior as an innate behavior is of prominent biological importance for survival and reproduction. However, molecular and circuit mechanisms underlying female sexual behavior is not well understood. Here, we identify the Cholecystokinin-like peptide Drosulfakinin (DSK) promotes female sexual behavior in *Drosophila*. Manipulation both *Dsk* and DSK neuronal activity impact female sexual receptivity. In addition, we reveal that *Dsk*-expressing neurons receive input signal from *R71G01GAL4* neurons to promote female sexual receptivity. Based on intersectional technique, we further found the regulation of female sexual behavior relies mainly on medial DSK neurons rather than lateral DSK neurons, and medial DSK neurons modulate female sexual behavior by acting on its receptor CCKLR-17D3. Thus, we characterized DSK/CCKLR-17D3 as *R71G01GAL4* neurons downstream signaling to regulate female sexual behavior.

## Introduction

In *Drosophila melanogaster*, innate sexual behavior is critical for reproduction and can induce several physiological and behavioral changes such as receptivity (Connolly and Cook, 1973), egg production (Bloch Qazi et al., 2003; Chen et al., 1988; Wang et al., 2020), female longevity (Chapman et al., 1995) and dietary preferences (Ribeiro and Dickson, 2010; Vargas et al., 2010). Thus, it is essential to understand and identify genetic and neural circuits that modulate innate sexual behavior. For male sexual behavior, a number of genes (Billeter et al., 2002; Emmons and Lipton, 2003) controlling male courtship have been identified and corresponding neural circuits have been dissected (Broughton et al., 2004; Clowney et al., 2015; Demir and Dickson, 2005; Kimura et al., 2008; Kohatsu et al., 2011; Pan and Baker, 2014; Ryner et al., 1996; Stockinger et al., 2005; Tanaka et al., 2017; Yamamoto and Koganezawa, 2013; Yu et al., 2010), whereas molecular and circuit mechanisms underlying female sexual behavior is less clear.

In recent years, genetic studies have shown that several genes play a critical role in regulating female sexual behavior. For example, mutant female of *spinster*, *icebox* and *chaste* show lower mating success rates while mutant female of *pain* show higher mating success rates than wild-type female (Carhan et al., 2005; Juni and Yamamoto, 2009; Kerr et al., 1997; Sakai et al., 2009; Suzuki et al., 1997). Moreover, specific subsets of neurons are required, both in the brain and ventral nerve cord, for modulating female sexual behavior. A significant decline of female sexual receptivity is observed when silencing specific neuron clusters in the central brain, such as two subsets of *doublesex*-expressing neurons (pCd and pC1) and two interneuron clusters (Spin-A and Spin-D) (Sakurai et al., 2013; Zhou et al., 2014). Female-specific vpoDNs in the brain integrate mating status and song to control both vaginal plate opening and female receptivity (Wang et al., 2021). Silencing either Abd-B neurons or SAG neurons located in the abdominal ganglion reduce female sexual receptivity (Bussell et al., 2014; Feng et al., 2014). In addition, female sexual behavior is also modulated by monoamines. In particular, dopamine not only plays a key role in regulating female sexual receptivity (Neckameyer, 1998), but also controls behavioral switching from rejection to acceptance in virgin females (Ishimoto and Kamikouchi, 2020); and octopamine is pivotal to female sexual behavior (Rezaval et al., 2014). The neuropeptides including SIFamide and Mip neuropeptide are responsible for female sexual receptivity (Jang et al.,2017; Terhzaz et al., 2007). Still, we know very little on how peptidergic neurons to control female sexual receptivity.

Drosulfakinin (DSK) is a neuropeptide, which is ortholog of Cholecystokinin (CCK) in mammals and its two receptors (CCKLR-17D1 and CCKLR-17D3) have been identified in *Drosophila* (Chen and Ganetzky, 2012; Kubiak et al., 2002; Nichols et al., 1988; Staljanssens et al., 2011), and previous studies have revealed that DSK peptide is involved in multifarious regulatory functions including satiety/food ingestion (Nassel and Williams, 2014; Williams et al., 2014), male courtship (Wu et al., 2019), aggression (Wu et al., 2020). However, it is not known whether DSK is crucial for female sexual behavior, and which specific neuron clusters that interact with DSK neurons to control female sexual behavior have not been identified.

In this study, 1) we reveal that DSK peptide act on its receptor CCKLR-17D3 to regulate female sexual behavior, 2) We found that *Dsk*-expressing neurons receive input from *R71G01GAL4* neurons and are required for female sexual behavior. Our results identify a neural subset of DSK neurons which function downstream of *R71G01GAL4* neurons and act through DSK receptor (CCKLR-17D3) neurons to control female sexual behavior.

## Results

### Dsk Gene Is Crucial for Female Receptivity in Virgin Female

To investigate the potential function of DSK peptide in modulating female sexual behavior, we first constructed knock out line for *Dsk* (*Figure 1A-D*) (Wu et al., 2020), and monitored the effect of *Dsk* mutant on female receptivity. Two parameters that copulation rate and latency were used to characterize receptivity (Ferveur, 2010). Interestingly, *Dsk* null mutant displayed reduced copulation rate and prolonged copulation latency compared with wild-type (*Figure 1E*) and heterozygous virgin females (*Figure 1F*). Moreover, RNAi knockdown of *Dsk* under the control of a pan-neuronal *elav^GAL4^* driver also reduced female receptivity (*Figure 1-Figure supplement 1A*). No significant change of locomotion activity was detected in *Dsk* mutant or RNAi-mediated females (*Figure 1-Figure supplement 2A-B*), and no significant difference was observed in courtship level in wild-type males paired with females of *Dsk* mutant and control (*Figure 1-Figure supplement 3A-B*). We note that manipulation of *Dsk* did not trigger egg-laying in virgin female and we also did not notice mating-induced TRIC signal changes (*Figure 1-Figure supplement 4A-B*).

**Figure 1.**
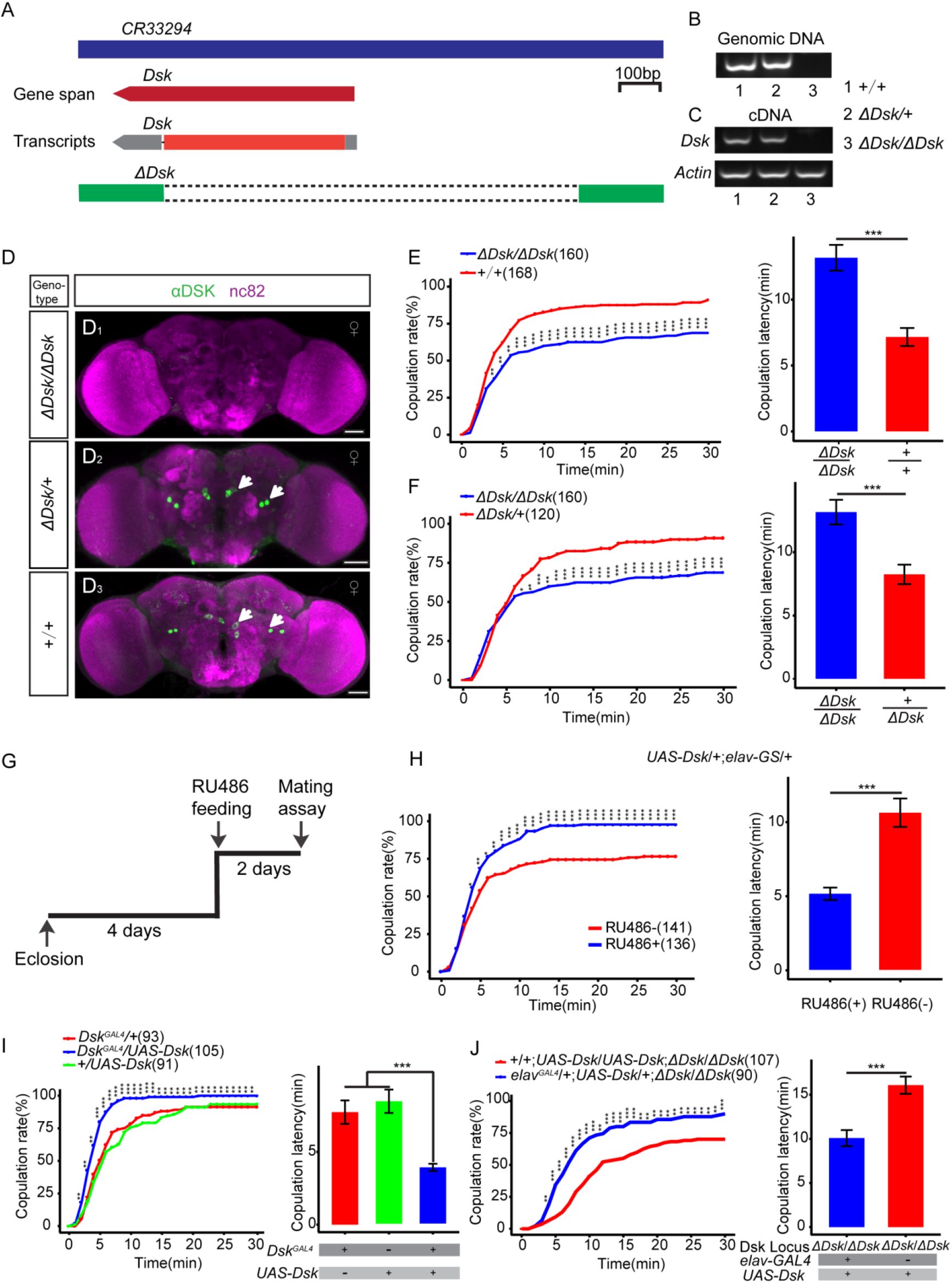
Dsk gene is important for female receptivity. (A) Organization of *Dsk* gene and generation of *ΔDsk*. (B-C) Validation of *ΔDsk*. RCR analysis from genomic DNA samples of *ΔDsk/ΔDsk*, *+/ΔDsk*, *+/+* (B), RT-PCR analysis from cDNA samples of *ΔDsk/ΔDsk*, *+/ΔDsk*, *+/+* (C). (D) Brain of indicated genotype, immunostained with anti-DSK antibody (green) and counterstained with nc82 (magenta). Arrow show cell bodies (green) stained with anti-DSK antibody. Scale bars: 50μm. (E-F) Receptivity of virgin females within 30min. *Dsk* mutant show reduced copulation rate and prolonged copulation latency compared with wild-type (E) and heterozygous females (F). (G) Schematic of experimental design. (H) Conditional overexpression of *Dsk* under the control of elav-GeneSwitch *(elav-GS)* significantly increased copulation rate and shortened copulation latency after feeding RU486 compared without feeding RU486. (I) Overexpression of *Dsk* in DSK neurons significantly increased copulation rate and shortened copulation latency compared with genetic controls. (J) Decreased female sexual behavior phenotypes of *ΔDsk/ΔDsk* were rescued by *elav^GAL4^* driving *UAS-Dsk*. The number of female flies paired with wild-type males analyzed is displayed in parentheses. And there are same numbers in two parameters. *p<0.05, **p<0.01, ***p<0.001, NS indicates no significant difference (chi-square test).

Next, we further investigated the importance of *Dsk* in controlling female sexual behavior. Conditional overexpression of *Dsk* under the control of elav-GeneSwitch (elav-GS), a RU486-dependent pan-neuronal driver (Osterwalder et al., 2001), induced copulation more quickly than control (*Figure 1G-H, Figure 1-Figure supplement 5A-B*). In addition, overexpression of *Dsk* in DSK neurons using *Dsk^GAL4^* to drive *UAS-Dsk* also increased copulation rate and shortened copulation latency compared with genetic controls (*Figure 2I*). Furthermore, we also carried out genetic rescue experiments to further confirm the importance of *Dsk* in modulating female sexual receptivity. To address this question, we used *elav^GAL4^*, a pan-neuronal driver, to drive *UAS-Dsk* expression in *Dsk* mutant background, and found that neuron-specific rescue with *UAS-Dsk* could restore the decreased receptivity level to normal level (*Figure 2J*). Taken together, these results indicated the function of *Dsk* is crucial for female sexual receptivity.

**Figure 2.**
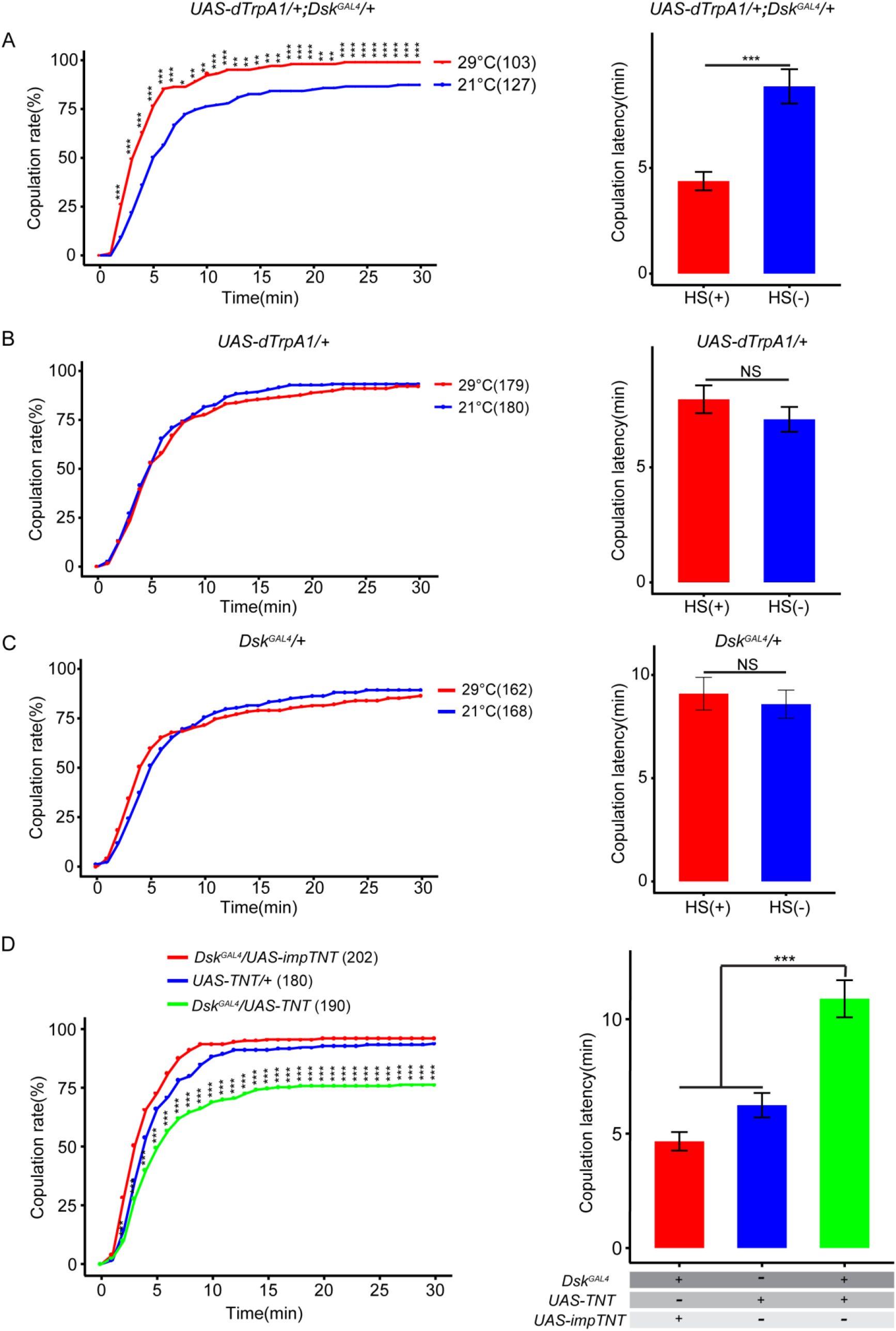
Effect of DSK neurons on female receptivity. (A) Activation of DSK neurons significantly increased copulation rate and shortened copulation latency at 29°C relative to 21°C. *Dsk^GAL^*^4^ driving *UAS-dTrpA1* activated DSK neurons at 29°C. (B-C) The controls with either *UAS-dTrpA1* alone or *Dsk^GAL^*^4^ alone did not alter the copulation rate and copulation latency at 29°C relative to 21°C. Activation of neuron is represented by (HS+) and control is represented by (HS-). (D) Inactivation of DSK neurons significantly decreased copulation rate and prolonged copulation latency compared with controls. *Dsk^GAL^*^4^ driving *UAS-TNT* inactivated DSK neurons. The number of female flies paired with wild-type males analyzed is displayed in parentheses. And there are same numbers in two parameters. *p<0.05, **p<0.01, ***p<0.001, NS indicates no significant difference (chi-square test).

### DSK Neurons Are Important for Modulating Female Receptivity

To further evaluate whether DSK neurons were involved in modulating female sexual behavior, we first activated DSK neurons by expressing the heat-activated *Drosophila* transient receptor potential channel (*dTrpA*) using a knock-in *Dsk^GAL4^* line (Hamada et al., 2008). Activation of DSK neurons increased female receptivity at 29°C relative to 21°C (*Figure 2A*). Whereas the female receptivity was not changed between 29°C and 21°C in controls with either *UAS-dTrpA1* alone or *Dsk^GAL4^* alone (*Figure 2B and 2C*). Activation of DSK neurons did not alter female receptivity in very young virgins and mated females (*TableS1*). Next, we use two strategies to inactivate DSK neurons by using *Dsk^GAL4^* to express either tetanus toxin light chain (TNT) which blocks synaptic vesicle exocytosis (Sweeney et al., 1995) or an inwardly rectifier potassium channel (Kir2.1) which hyperpolarizes neurons and suppress neural activity (Baines et al., 2001; Thum et al., 2006). Female receptivity was decreased when silencing DSK neurons either using TNT (*Figure 2D*) or using Kir2.1 (*Figure 2-Figure supplement 1A*). Thus, our results suggest that DSK neurons are important for modulating female sexual behavior.

### Anatomical and Functional Dissection of the Connection Between DSK Neurons and R71G01GAL4 Neurons

Recent studies revealed that DSK neurons not only function downstream of a subset of P1 neurons to regulate aggressive behavior (Wu et al., 2020), but also function antagonistically with P1 neurons to regulate male courtship (Wu et al., 2019) in males. It was known that *R71G01GAL4* neurons also expressed in female brains (Hoopfer et al., 2015), and activation of *R71G01GAL4* neurons promoted female receptivity (*Figure 3-supplement 1A*). Given that, we hypothesized whether DSK neurons are functional targets of *R71G01GAL4* neurons in controlling female receptivity. Thus, we first sought to detect whether *Dsk*-expressing neurons had potential synaptic connection with *R71G01GAL4* neurons via GRASP method (GFP reconstitution across synaptic partners) (Feinberg et al., 2008; Gordon and Scott, 2009). Interestingly, we detected significant recombinant GFP signals (GRASP) between *R71G01GAL4* neurons and DSK neurons (*Figure 3-supplement 2A-C*), suggesting that these neurons form a direct synaptic connection. Next, we used *trans*-Tango (Talay et al., 2017) approach to further confirm whether *Dsk*-expressing neurons are immediate downstream of *R71G01GAL4* neurons. Interestingly, *trans*-Tango signal was observed in DSK neurons by co-staining the *trans*-Tango flies with DSK antibodies (*Figure 3A-B, Figure 3-supplement 3A*). Moreover, we registrated *R71G01GAL4* neurons and DSK neurons, and found that *R71G01GAL4* neurons axons overlap with DSK neurons dendrites (*Figure 3C*).

**Figure 3.**
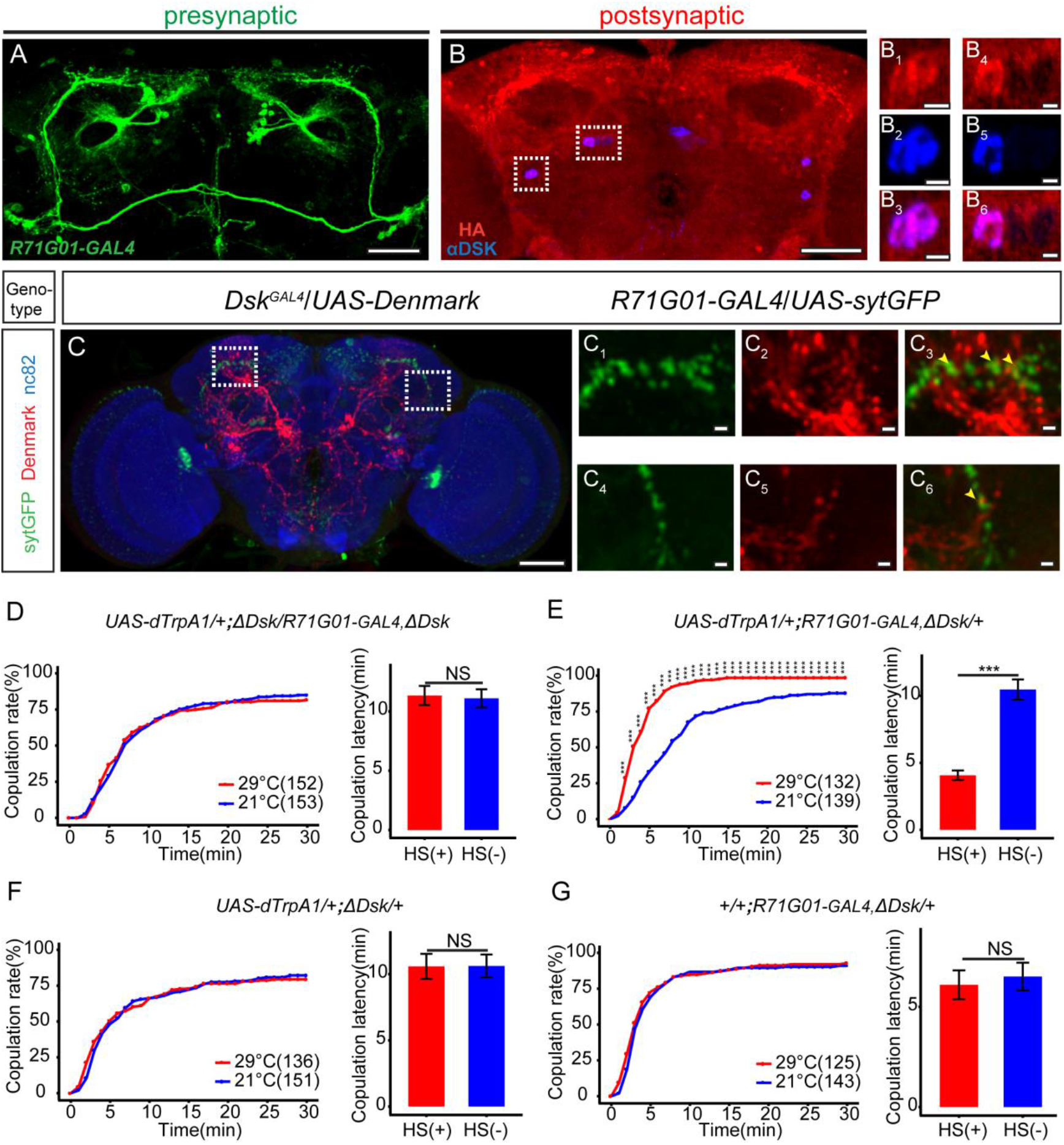
DSK neurons are functional targets of R71G01GAL4 neurons in regulating mating behavior. (A-B) Transsynaptic circuit analysis using *trans*-Tango confirms that *Dsk*-expressing neurons are postsynaptic neurons of *R71G01GAL4* neurons. In the central brain, expression of the Tango ligand in *R71G01GAL4* neurons (green) (A) induced postsynaptic mtdTomato signals (anti-HA, red) (B). Cell bodies of Dsk were stained with anti-DSK (blue) (B). Magnification of white boxed region in (B) is shown in (B_1_-B_3_) and (B_4_-B_6_). Scale bars are 50μm in (A-B), 5μm in (B_1_-B_3_) and (B_4_-B_6_). (C) *R71G01GAL4* neurons axons overlapped with DSK neurons dendrites by anatomical registration. Magnification of white boxed region in (C) is shown in (C_1_-C_3_) and (C_4_-C_6_). Yellow arrowheads indicated the region of overlaps between *R71G01GAL4* neurons axons with DSK neurons dendrites. *R71G01-GAL4* driven *UAS-sytGFP* expression (green), *Dsk^GAL4^* driven *UAS-Denmark* expression (red). Scale bars are 50μm in (C), 5μm in (C_1_-C_3_) and (C_4_-C_6_). (D) The copulation rate and copulation latency had no difference at 29°C relative to 21°C in the case of activation of *R71G01GAL4* neurons in the *ΔDsk* mutant background. (E) The positive control significantly increased copulation rate and shortened copulation latency at 29°C relative to 21°C. (F-G) The negative controls did not alter the copulation rate and copulation latency by heating. Activation of neuron is represented by (HS+) and control is represented by (HS-). The number of female flies paired with wild-type males analyzed is displayed in parentheses. And there are same numbers in two parameters. *p<0.05, **p<0.01, ***p<0.001, NS indicates no significant difference (chi-square test).

In addition, we performed behavioral epistasis experiment to confirm functional interactions between DSK neurons and *R71G01GAL4* neurons. We activated *R71G01GAL4* neurons by *R71G01-GAL4* driving *UAS-dTRPA1* in the *Dsk* mutant background, and found that increased levels of female receptivity caused by activation of *R71G01GAL4* neurons were suppressed by the mutation in *Dsk* (*Figure 3D-G*). Taken together, these results further demonstrate that DSK neurons are the functional targets of *R71G01GAL4* neurons in controlling female sexual behavior.

### Medial DSK Neurons (DSK-M) Rather Than Lateral DSK Neurons (DSK-L) Are Critical for Regulating Female Receptivity

Previous study has implicated that eight DSK neurons were classified two types (DSK-M and DSK-L) based on the location of the cell bodies in female and male brains (Wu et al., 2020). However, functional difference between those two types was not characterized in female files.

We first performed patch-clamp recordings on DSK-M and DSK-L neurons, respectively, to confirm the functional connectivity between *R71G01GAL4* neurons and *Dsk*-expressing neurons. We activated *R71G01GAL4* neurons through ATP activation of ATP gating of P2X_2_ transgenic receptors (Brake et al., 1994; Macara et al., 2012) and recorded the electrical responses in DSK-M neurons and DSK-L neurons. In perforate patch recordings, ATP/P2X_2_ activation of *R71G01GAL4* neurons induced strong electrical responses from DSK-M neurons and relatively weak responses from DSK-L neurons in female (*Figure 4A-C*).

**Figure 4.**
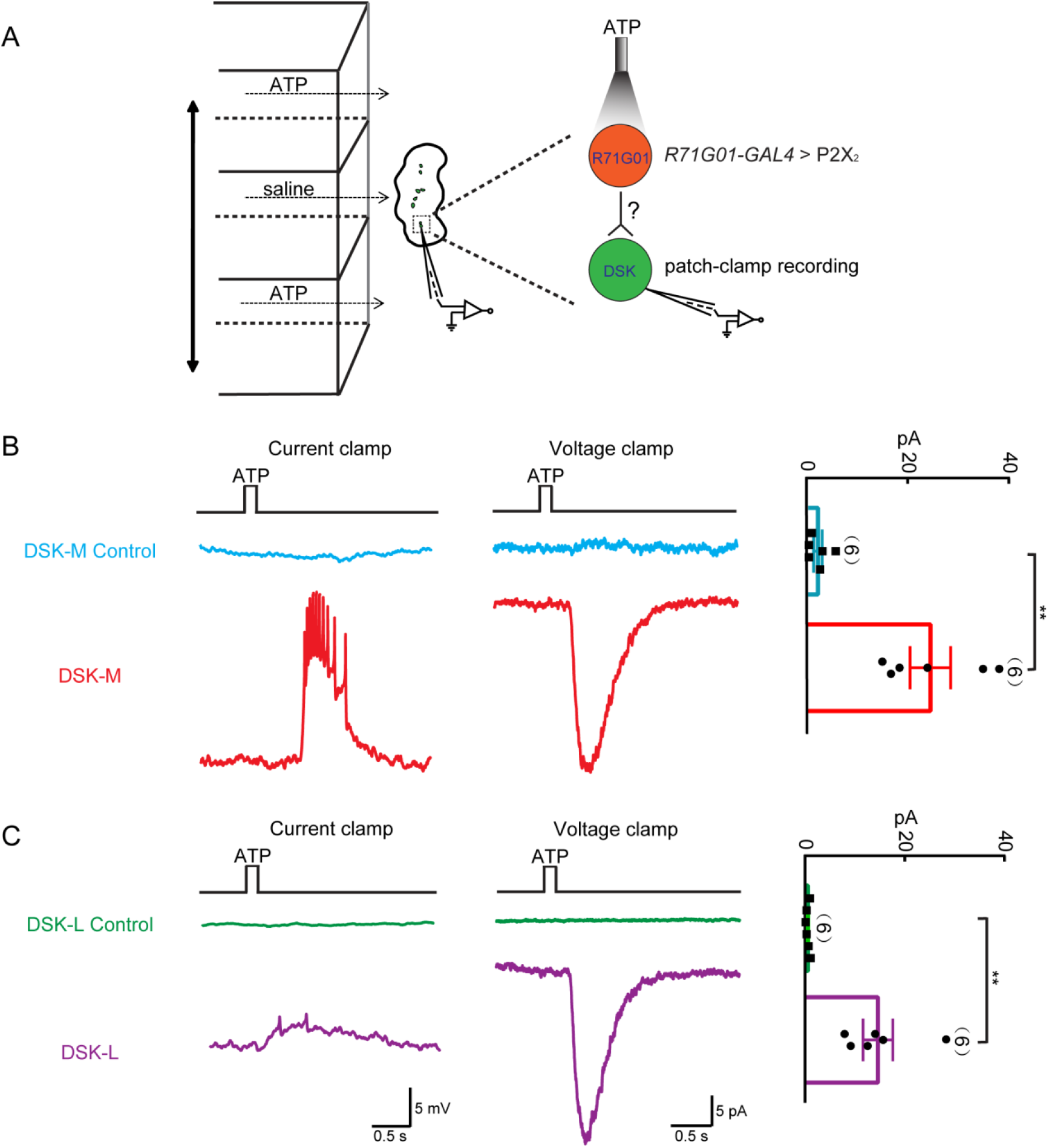
Functional connectivity between R71G01GAL4 neurons and DSK neurons. (A) Schematic illustration of activation of *R71G01GAL4* neurons by ATP and patch-camp recording of DSK neurons. *R71G01GAL4* neurons were activated by ATP from *+/+;R71G01-LexA/+;Dsk^GAL4^/LexAop-P2X_2_,UAS-GCaMP6m* files. (B-C) The electrical responses of medial DSK neurons (DSK-M) and lateral DSK neurons (DSK-L) to the ATP activation of P2X_2_-expressing *R71G01GAL4* neurons. ATP: 2.5 mM. Left: ATP-induced spiking firing (current clamp). Middle: current responses (voltage clamp). Right: quantification of absolute current responses. n=6 for DSK-M, DSK-M control, DSK-L, DSK-L control. Genotype: *+/+;+/+;Dsk^GAL4^/LexAop-P2X_2_,UAS-GCaMP6m* for DSK-M control and DSK-L control. **p<0.01 (Mann-Whitney U tests).

To further distinguish if both types are or only one type of DSK neurons was involved in regulating female sexual behavior, we used intersectional strategy to subdivide DSK neurons and manipulated DSK-M and DSK-L separately. We first screened about 100 knock-in lines containing GAL4 of *Drosophila* chemoconnectome (CCT) (Deng et al., 2019) combined with *DskFlp* line to drive *UAS>stop>myr*∷*GFP* (a Gal4/Flp-responsive membrane reporter) expression, and further identified the GFP signal using anti-DSK antibodies. Interestingly, we found intersection of *DskFlp* and *GluRIA^GAL4^* specifically labels DSK-M neurons (*Figure 5A*), and intersection of *DskFlp* and *TβH^GAL4^* specifically labels DSK-L neurons (*Figure 5B*). Next, we confirmed the behavioral relevance of specific subtypes of DSK neurons. Activation of DSK-M neurons significantly increased female receptivity (*Figure 5C, Figure 5-supplement 1A-C*) while inactivation of DSK-M neurons significantly reduced female receptivity, neither of these effects was observed in genetic controls (*Figure 5D*). Thus, DSK-M neurons are responsible for regulating female sexual receptivity. Conversely, neither activation nor inactivation of DSK-L neurons altered the female receptivity level in virgin females (*Figure 5E-F, Figure 5-supplement 1C-E*). Taken together, these findings indicate that DSK-M neurons, rather than DSK-L neurons, play an essential role in regulating female sexual behavior.

**Figure 5.**
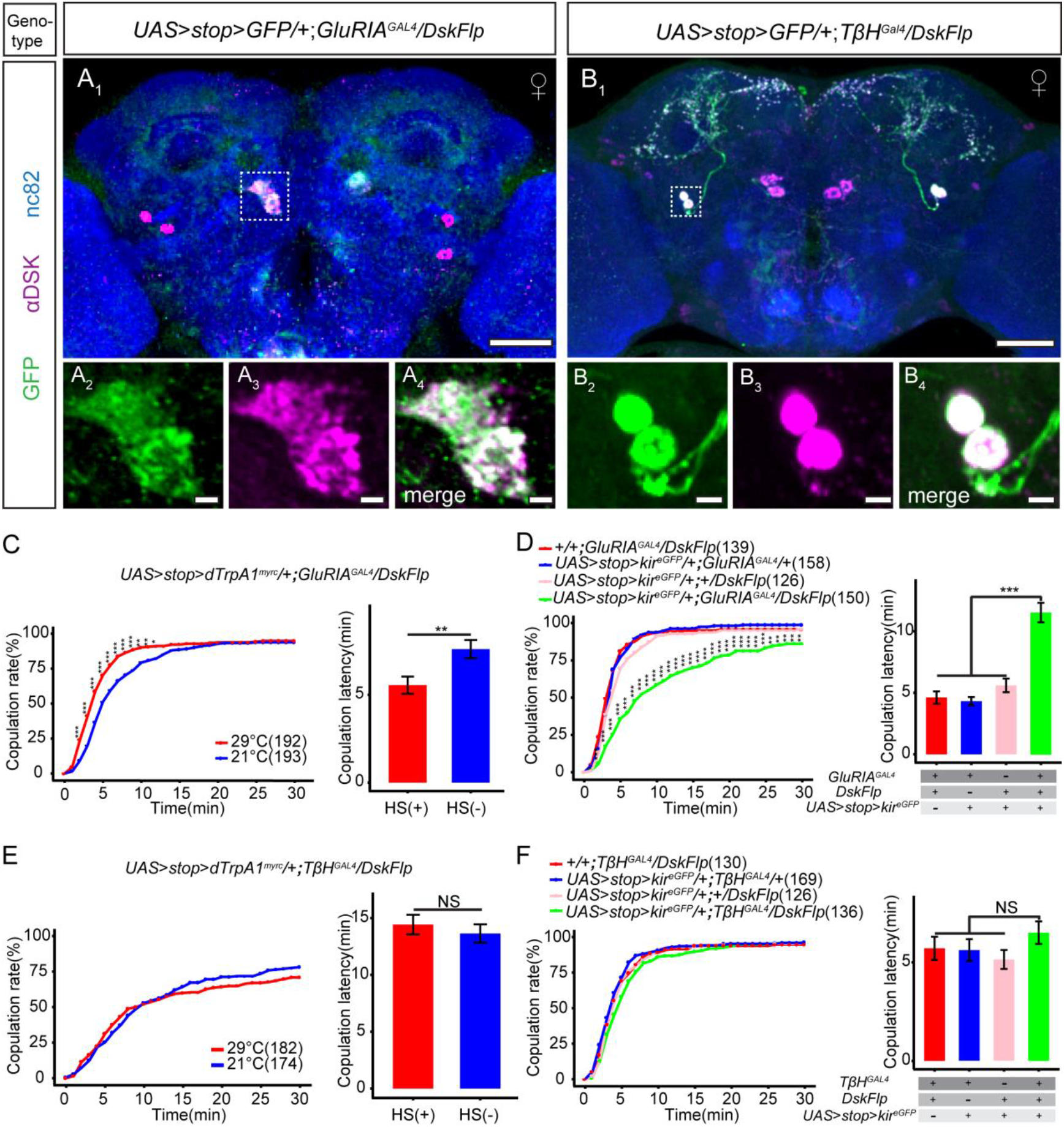
DSK-M neurons play a critical role in female receptivity. (A) Intersectional expression of *Dsk* neurons and *GluRIA* neurons were detected by immunostained with anti-GFP (green) and anti-DSK (magenta) antibodies in female brain and were counterstained with anti-nc82 (blue). Magnification of white boxed region in (A) is shown in (A_2_-A_4_). Genotype: *UAS>stop>myr*∷*GFP*/+*;GluRIA^GAL4^/DskFlp.* (B) Intersectional expression of *Dsk* neurons and *TβH* neurons were detected by immunostained with anti-GFP (green) and anti-DSK (magenta) antibodies in female brain and were counterstained with anti-nc82 (blue). Magnification of white boxed region in (B) is shown in (B_2_-B_4_). Genotype: *UAS>stop>myr*∷*GFP*/+*;TβH^GAL4^/DskFlp.* Scale bars are 50μm in (A and B), 5μm in (A_1_-A_3_) and (B_1_-B_3_). (C) Activation of co-expression neurons of *Dsk* and *GluRIA* significantly increased copulation rate and shortened copulation latency at 29°C relative to 21°C. Genotype: *UAS>stop>dTrpA^myrc^*/+*;GluRIA^GAL4^/DskFlp.* (D) Inactivation of co-expression neurons of *Dsk* and *GluRIA* significantly decreased the copulation rate and prolonged copulation latency compared with controls. Genotype: *UAS>stop>kir^eGFP^*/+*;GluRIA^GAL4^/DskFlp*, +/+*;GluRIA^GAL4^/DskFlp, UAS>stop>kir^eGFP^*/+*; GluRIA^GAL4^*/+, *UAS>stop>kir^eGFP^*/+;+/*DskFlp.* (E) Activation of co-expression neurons of *Dsk* and *TβH* did not alter the copulation rate and copulation latency at 29°C relative to 21°C. Genotype: *UAS>stop>dTrpA^myrc^*/+*;TβH^GAL4^/DskFlp.* (F) Inactivation of co-expression neurons of *Dsk* and *TβH* did not alter the copulation rate and copulation latency compared with controls. Genotype: *UAS>stop>kir^eGFP^*/+*;TβH^GAL4^/DskFlp*, *UAS>stop>kir^eGFP^*/+;+/*DskFlp*, *UAS>stop>kir^eGFP^*/+*;TβH^GAL4^*/+, +/+*;TβH^GAL4^/DskFlp*. Activation of neuron is represented by (HS+) and control is represented by (HS-). The number of female flies paired with wild-type males analyzed is displayed in parentheses. And there are same numbers in two parameters. *p<0.05, **p<0.01, ***p<0.001, NS indicates no significant difference (chi-square test).

### *Dsk* Regulates Female Receptivity via Its Receptor *CCKLR-17D3*

Next, we want to explore the downstream target of DSK neurons. Two *Dsk* receptors were identified: *CCKLR-17D1* and *CCKLR-17D3* (Chen and Ganetzky, 2012; Kubiak et al., 2002), and it would be essential to distinguish which receptor is or both of receptors are critical for modulating female sexual behavior. We constructed knock out lines for these two receptors (*Figure 6A-C*) (Wu et al., 2020). Knock out of *CCKLR-17D3* rather than *CCKLR-17D1* reduced mating success rate in virgin females compared with control flies (*Figure 6D, Figure 6-supplement 1A*). Moreover, RNAi knockdown of *CCKLR-17D3* under the control of a pan-neuronal *elav^GAL4^* driver or *CCKLR-17D3^GAL4^* also significantly reduced female receptivity (*Figure 6E, Figure 6-suplement 2A*). Conditional silencing of *CCKLR-17D3* using the elav-GeneSwitch (elav-GS) system to avert the effect of the genetic background also significantly decreased female receptivity (*Figure 6-suplement 2B-D*). In addition, no significant change of locomotion activity was detected in *CCKLR-17D3* mutant or silencing of *CCKLR-17D3* (*Figure 6-supplement 3A-B*). The reduced female receptivity phenotypes of *ΔCCKLR-17D3/ΔCCKLR-17D3* could be rescued by expression of *UAS-CCKLR-17D3* driven by *elav-GS* (*Figure 6F-H*). These results demonstrate that *CCKLR-17D3* is critical for modulating female sexual receptivity.

**Figure 6.**
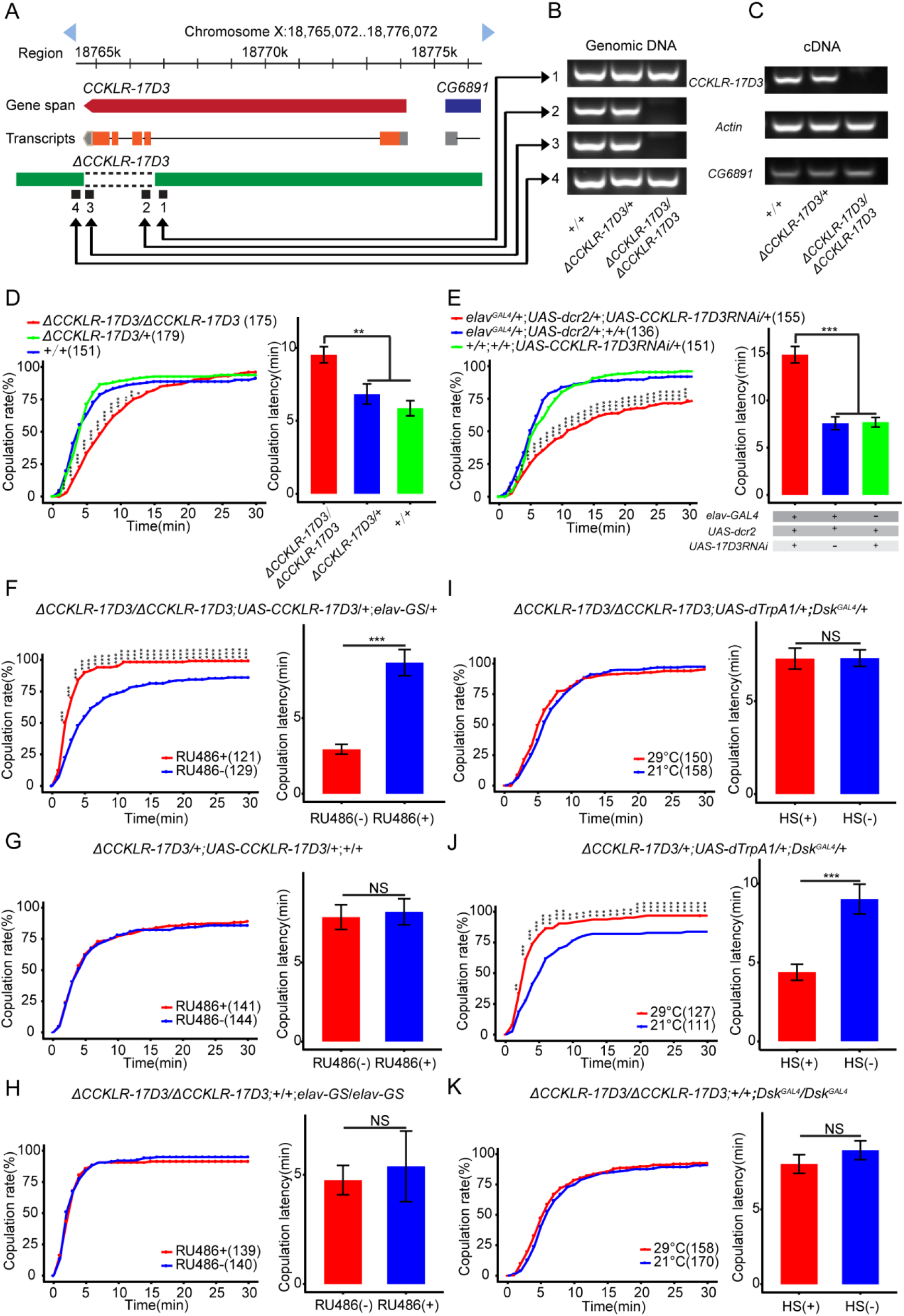
*Dsk* regulate female receptivity via *CCKLR-17D3* Receptor. (A) Organization of *CCKLR-17D3* and generation of *ΔCCKLR-17D3*. (B-C) Validation of *ΔCCKLR-17D3*. RCR analysis from genomic DNA samples of *ΔCCKLR-17D3/ΔCCKLR-17D3*, *+/ΔCCKLR-17D3*, *+/+*. (B) RT-PCR analysis from cDNA samples of *ΔCCKLR-17D3/ΔCCKLR-17D3*, *+/ΔCCKLR-17D3*, *+/+* (C). (D) *CCKLR-17D3* mutant females significantly decreased copulation rate and prolonged copulation latency compared with wild-type and heterozygous. (E) Knockdown of *CCKLR-17D3* showed significantly reduced copulation rate and prolonged copulation latency compared with controls. *CCKLR-17D3*RNAi was driven by *elav-Gal4;UAS-dcr2*. (F) Conditional expression of *UAS-CCKLR-17D3* in the *ΔCCKLR-17D3* mutant background after feeding RU486 significantly increased copulation rate and shortened copulation latency compared without feeding RU486. (G-H) The controls with either *UAS-CCKLR-17D3* alone or *elav-GS* alone did not recue the phenotypes of *ΔCCKLR-17D3/ΔCCKLR-17D3* at feeding RU486 relative to without feeding RU486. (I) The copulation rate and copulation latency have no difference at 29°C relative to 21°C in the case of activating DSK neurons in the *ΔCCKLR-17D3* mutant background. (J) The positive control significantly increased copulation rate and shortened copulation latency at 29°C relative to 21°C. (K) The negative control did not alter the copulation rate and copulation latency by heating. Activation of neuron is represented by (HS+) and control is represented by (HS-). The number of female flies paired with wild-type males analyzed is displayed in parentheses. And there are same numbers in two parameters. *p<0.05, **p<0.01, ***p<0.001, NS indicates no significant difference (chi-square test).

To ask whether the enhanced receptivity level of DSK neuronal activation in females relied on downstream CCKLR-17D3 or not, we studied the female receptivity of *CCKLR-17D3* mutant flies with DSK neurons activated by TrpA1 using *Dsk^GAL4^* to drive *UAS-dTrpA1*. We found that knock-out of *CCKLR-17D3* could block the increased levels of female receptivity caused by activating of DSK neurons (*Figure 6I-K*). These data support that DSK neurons rely on its receptor CCKLR-17D3 to regulate female sexual receptivity.

## Discussion

In this study, we systematically investigated *Dsk*-mediated neuromodulation of female sexual receptivity. At the neuronal circuit level, we identified that DSK neurons are the immediate downstream targets of *R71G01GAL4* neurons in controlling female sexual receptivity. Moreover, we employ intersectional tool to subdivide DSK neurons into medial DSK neurons (DSK-M) and lateral DSK neurons (DSK-L) and uncover that DSK-M neurons rather than DSK-L neurons play essential roles in modulating female sexual receptivity. At the molecular level, we reveal that DSK neuropeptide and its receptor CCKLR-17D3 are critical for modulating female sexual receptivity. Collectively, our findings illuminate that *R71G01GAL4*-DSK-M-CCKLR-17D3 signal forms a crucial pathway in regulating female sexual behaviors

The female sexual behavior is a complex innate behavior. Virgin female receives multiple sensory cues to decide whether or not to accept a courting male. pC1 neurons are required for modulating female sexual receptivity and this neural cluster integrates pheromone and song cues (Zhou et al., 2015; Zhou et al., 2014) and *R71G01GAL4* neurons may contain part of pC1 neurons. Our results not only demonstrate *Dsk* plays a key role in the regulation of female sexual receptivity, but also uncover DSK neurons are functional targets of *R71G01GAL4* neurons in regulating female sexual behavior. Thus, we would venture to speculate that DSK neurons could also integrate sensory stimuli like song or pheromone cues. If this speculation is true, it would be interesting to further investigate whether DSK neurons function downstream of pC1 neurons to control female receptivity, and how pC1 neurons and DSK neurons coordinate to integrate sensory stimuli to modulate female sexual behavior.

Whether or not a male is successful in courting to a female depends on female’s sexual maturity or mating status. Very young virgins display low receptivity level to courting males, mature virgin females show high receptivity level and are willing to accept courting males, and mated females become temporarily sexually unreceptive to courting males (Dickson, 2008; Kubli, 2003; Rezaval et al., 2012). In this study, we found that both *Dsk* and DSK neurons are involved in regulating virgin female sexual receptivity, so we asked whether manipulating DSK neurons could also affect female sexual receptivity in very young virgins or mated females. Activation of DSK neurons did not alter female receptivity in both very young virgins and mated females (*TableS1*). Thus, *Dsk* only plays a pivotal role in virgin females.

In the process of sexual behavior, if the female is willing to accept the male’s courtship, the female slow down their movement to allow the male to engage in copulation (Bussell et al., 2014; Tompkins et al., 1982). Hence, it is quite possible that decreased copulation rate of *Dsk*-ablated virgin females is due to the defect of locomotion activity. Conversely, the locomotion of *Dsk*-ablated female using Ctrax software (Branson et al., 2009) shows no difference compared with control females (*Figure 1-supplement 2A*), suggesting that less receptivity of *Dsk*-ablated virgin females is not caused by the change of locomotion activity. Reducing copulation rate in *Dsk*-ablated females may also be due to the abatement of female sexual appeal or the changes of male ardor to courtship. Nevertheless, no matter the object of virgin female targets be *ΔDsk/ΔDsk*, *ΔDsk/+*, *+/+*, the courting male display normal courtship level (*Figure 1-supplement 3A-B*). Taken together, the slow copulation in *Dsk*-ablated virgin female is attributed to decreased sexual receptivity.

Previous study has revealed that eight DSK neurons are classified two types (DSK-M and DSK-L) based on the location of the cell bodies, and DSK-M neurons extend descending fibers to ventral nerve cord (Wu et al., 2020). In this study, we also found that activating DSK-M neurons enhance female receptivity whereas inactivating DSK-M neurons reduce female receptivity. Silencing adult Abd-B neurons and SAG neurons located in the abdominal ganglion inhibits female sexual receptivity (Bussell et al., 2014; Feng et al., 2014). However, whether *Dsk* system connects with SAG neurons or Abd-B neurons in regulating female sexual behavior remain unknown and it’s interesting to further investigate this possibility. In addition, Wu and colleagues have reported that eight DSK neurons could be further classified into three neural types (Type I, II and III) based on the morphology of the neuritis, and Type I and II neurons correspond to DSK-M while Type III neurons correspond to DSK-L (Wu et al., 2020). DSK neurons modulate not only male courtship (Wu et al., 2019) but also male aggression (Wu et al., 2020). In this study, DSK neurons also modulate female sexual behavior. Thus, it is necessary to label neuronal type more specifically and identify whether different subtype involves in different behaviors, such as male courtship, aggression and female sexual behavior.

## EXPERIMENTAL PROCEDURES

### Fly stocks

Flies were reared on standard cornmeal-yeast medium under a 12 hr:12 hr dark:light cycle at 25°C and 60% humidity. Flies carrying a *dTrpA* transgene were raised at 21°C. *UAS-TNTE* and *UAS-impTNT* were kindly provided by Dr. Cahir O’Kane (University of Cambridge). *UAS-dTRPA1* was a gift from Dr. Paul Garrity (Brandeis University). *trans*-Tango line and *elav-GS* line were provided by Dr. Yi Zhong (Tsinghua University), *DskFlp* was provided by Dr. Yufeng Pan. *UAS>stop>myr∷GFP* (pJFRC41 in attP5) was a gift from Gerald Rubin, *UAS>stop>kir^eGFP^* was provided by Dr. Yi Rao. The following lines were obtained from the Bloomington *Drosophila* Stock Center: *R71G01-GAL4* (BL#39599), *R71G01-LexA* (BL#54733), *TβH-GAL4* (BL#45904), TRIC line (BL#61679), *UAS-Kir2.1* (BL#6595 and BL#6596), *UAS-mCD8∷GFP* (BL#5137), *UAS>stop>dTrpA^myrc^* (BL#66871). *Lexo-CD4-spGFP11/CyO*; *UAS-CD4-spGFP1-10/Tb* was previously described (Gordon and Scott, 2009).

### Method detail

#### Behavioral Assays

Flies were reared at 25°C. Virgin females and wild-type males were collected upon eclosion, placed in groups of 12 flies each and aged for 5-7d at 25°C and 60% humidity before carrying out behavior assay except the thermogenetic experiments.

In female sexual behavior experiment in virgin female, mating behavior assays were carried out in the courtship chamber. A virgin female of defined genotype and a wild-type male were gently cold anaesthetized and respectively introduced into two layers of the round courtship chambers in which separated by a removable transparent film. The flies were allowed to recover for at least 1h before the film was removed to allow the pair of a test female and a wild-type male to contact. The mating behavior was recorded using a camera (Canon VIXIA HF R500) for 30 min at 30 fps for further analysis.

For female sexual behavior experiment in very young virgin female, we collected flies with 0-3h post-eclosion and measures receptivity at 12-16h post-eclosion using same method as mentioned above.

For female sexual behavior experiment in mated female, we first collected virgin female upon eclosion and generate mated females by pairing females aged 5-7d with wild-type males. Mated females were isolated 18-24h and then assayed for receptivity with a new wild-type male using same method as mentioned above.

For *dTrpA* activation experiment, flies were reared at 21°C. Flies were loaded into courtship chamber and recovered at least 30min, then were placed at 21°C (control group) or 29°C (experimental group) for 30min before removing the film and videotaping.

For egg laying experiment, virgin females were collected upon eclosion and 5 flies were housed on standard medium in single vials. The flies were transferred into new food tubes every 24h, and we manually counted the number of eggs in each food tube.

#### Immunohistochemistry

Whole brains of flies aged for 5-7d were dissected in 1X PBS and fixed in 2% paraformaldehyde for 55 min at room temperature. The samples were blocked in 5% normal goat serum for 1 hr at room temperature after washing the samples with PBT (1X PBS containing 0.3% Triton-X100) for four times for 15 min. Then the samples were incubated in primary antibodies (diluted in blocking solution) for 18-24hr at 4°C. Samples were washed four times with 0.3% PBT for 15 min, then were incubated in secondary antibodies (diluted in blocking solution) for 18-24hr at 4°C. Samples were washed four times with 0.3% PBT for 15 min, then were fixed in 4% paraformaldehyde for 4hr at room temperature. Finally, brains were mounted on poly-L-lysine (PLL)-coated coverslip in 1X PBS. The coverslip were dipped for 5min with ethanol of 30%->50%->70%->95%->100% sequentially at room temperature, and then dipped three times for 5min with Xylene. The brains were mounted with DPX and allowed DPX to dry for 2 days before imaging. Confocal images were obtained with Carl Zeiss (LSM710) confocal microscopes and Fiji software was used to process images. Primary antibodies used were: chicken anti-GFP (1:1000; life technologies), rabbit anti-DSK antibody (1:1000), mouse anti-nc82 (1:50; DSHB), Rat anti-HA (1:100; Roche), mouse anti-GFP-20 (1:100; sigma). Secondary antibodies used were: Alexa Fluor goat anti-chicken 488 (1:500; life technologies), Alexa Fluor goat anti-rabbit 546 (1:500; life technologies), Alexa Fluor goat anti-mouse 647 (1:500; life technologies), Alexa Fluor goat anti-rat 546 (1:500; Invitrogen) and Alexa Fluor goat anti-mouse 488 (1:500; life technologies).

#### Generation of *UAS-CCKLR-17D3*

The method of generation of *UAS-CCKLR-17D3* was same as described previously (Wu et al., 2020). Primer sequences for cloning the cDNA of *UAS-CCKLR-17D3* are as follows: *UAS-CCKLR-17D3*:

Forward:

ATTCTTATCCTTTACTTCAGGCGGCCGCAAAATGTTCAACTACGAGGAGGG

Reverse:

GTTATTTTAAAAACGATTCATTCTAGATTAGAGCTGAGGACTGTTGACG

#### Genomic DNA extraction and RT-RCR

Genomic DNA was extracted from whole fly body using MightyPrep reagent for DNA (Takara). Whole head RNA was extracted from 50 fly heads using TRIzol (Ambion #15596018). cDNA was generated from total RNA using the Prime Script reagent kit (Takara).

#### Validation of Δ*CCKLR-17D3* in female

Candidates of Δ*CCKLR-17D3* were characterized by the loss of DNA band in the deleted areas by PCR on the genomic DNA, as shown in Figure 6A. Primer sequences used for regions 1–4 in Figure 6B are as follows:

Region (1): Forward 5’-CAGTAGAGGATTCGCCTCCAAG-3’

Reverse 5’-GACATACAGCGAGAGTGC-3’
Region (2): Forward 5’-CATGAACGCCAGCTTCCG-3’

Reverse 5’-GCACTATTGGTGGTCACCAC-3’
Region (3): Forward 5’-GGAAATCATCTAACAGGCTTAC-3’

Reverse 5’-GCCGTGTCAAATCGCTTTC-3’
Region (4): Forward 5’-GCATACATACAAGCAAATTATGC-3’

Reverse 5’-CTCATATTCTTTTGGGCTACCAC-3’

Primer sequences used for amplifying *CCKLR-17D3* or *CG6891* cDNA in Figure 6C are as follows:

*CCKLR-17D3* cDNA: Forward 5’-GCCCATAGCGGTCTTTAGTC-3’

Reverse 5’-GTGATGAGGATGTAGGCCAC 3
*CG6891* cDNA: Forward 5’-GCTGTGTTCTGGATGTGGATG-3’

Reverse 5’-CTGGAACTGTGCTGGTTCTG-3’

#### Drug Feeding

Virgin females of defined genotype were collected upon eclosion and reared on standard cornmeal-yeast medium as a group of 12 for 4 days. Then we transferred the female flies to new standard cornmeal-yeast food tube containing 500μM RU486 (RU486+) or control solution (RU486-) for 2 days before behavior assay. RU486 (mifepristone; Sigma) was dissolved in ethanol.

#### TRIC analysis

*Dsk^GAL4^* flies were crossed with a TRIC line to detect the changes of intracellular Ca^2+^ levels between virgin and mated females. The brain of virgin and mated female (2 days after copulation) were dissected and fixed with 8% paraformaldehyde for 2hr, and then mounted with DPX. All of the confocal images were obtained with Carl Zeiss (LSM710) confocal microscopes with the same settings.

Fiji software was used to process images. We first generated a Z stack of the sum of fluorescence signals, and then quantified the fluorescence intensity of DSK cell bodies between virgin and mated female brain, respectively. We calculated the signal intensity of each cell by setting the average of the cell in each group from virgin females as 100%

#### Electrophysiological recordings

Young adult flies (1–2 days after eclosion) were anesthetized on ice and brain were dissected in saline solution. And the brain was continuously perfused with saline bubbled with 95% O_2_/5% CO_2_ (~pH 7.3) at room temperature. The saline composed the following (in mM): 103 NaCl, 3 KCl, 4 MgCl_2_, 1.5 CaCl_2_, 26 NaHCO_3_, 1 NaH_2_PO_4_, 5 N-tri-(hydroxymethyl)-methyl-2-aminoethane-sulfonic acid (TES), 20 D-glucose, 17 sucrose, and 5 trehalose.

Electrophysiological recordings were performed using a Nikon microscope with a 60 water immersion objective to locate target neurons. Then we used Nikon A1R+ confocal microscope with infrared-differential interference contrast (IR-DIC) optics to visual for patch-clamp recordings and the image was shown on monitor by IR-CCD (DAGE-MTI). The recording pipette (~10–15 MΩ) was filled with internal solution containing 150 mg/ml amphotericin B. The internal solution consists of (in mM): 140 K-gluconate, 6 NaCl, 2 MgCl2, 0.1 CaCl2, 1 EGTA, 10 HEPES (pH 7.3). Current and voltage signals were amplified with MultiClamp 700B, digitized with Digidata 1440A, recorded with Clampex 10.6 (all from Molecular Devices), filtered at 2 kHz, and sampled at 5 kHz. The recorded neuron was voltage clamped at 70 mV. Measured voltages were corrected for a liquid junction potential.

#### Chemogenetic stimulation

ATP-gated ion channel P2X_2_ was driven by *71G01-GAL4*. ATP-Na (Sigma-Aldrich) of 2.5 mM was delivered through a three-barrel tube, controlled by stepper (SF77B, Warner Instruments) driven by Axon Digidata 1440A analog voltage output. The aim of these equipments was for fast solution change between perfusion saline and ATP stimulation.

#### Brain image registration

A standard brain was generated using CMTK software as described previously (Rohlfing and Maurer, 2003; Zhou et al., 2014). Confocal stacks were then registered into the common standard brain with a Fiji graphical user interface (GUI) as described previously (Jefferis et al., 2007).

#### Quantification and statistical analysis of female mating behavior

Two parameters that copulation rate and latency were used to characterize receptivity. The time from removing the film to copulation was measured for each female. The number of females that had engaged in copulation by the end of each 1-min interval were summed within 30min and plotted as a percentage of total females for each time point. The time from removing the film to successful copulation for each female was used to characterize latency to copulation. And all the time point that female successfully copulated were analyzed by manual method and unhealthy flies were discarded. Three scorers with blinding to the genotypes and condition of the experiment were assigned for independent scoring.

## Statistical analysis

Statistical analyses were carried about using R software version 3.4.3 or GraphPad software. Chi-square tests were used for comparing different groups in the female receptivity assay. For locomotion or male courtship experiments, Kruskal-Wallis ANOVA test followed by post-hoc Mann-Whitney U test was used for comparison among multiple groups. The Mann-Whitney U test was applied for analyzing the significance of two columns.

## Acknowledgements

We thank Yi Rao (Peking University), Yi Zhong (Tsinghua University), Yufeng Pan (Southeast University), Cahir O’Kane (University of Cambridge) and Paul Garrity (Brandeis University) for providing fly lines. We thank Pengxiang Wu (Chinese Academy of Sciences) and Yu Mu (Institute of neuroscience) for comments on the manuscript; Yihui Chen for the maintenance of fly stocks; other members of the Zhou laboratory for helpful discussion. This work is supported by grants to Chuan Zhou from National Natural Science Foundation of China (NO.Y711241133) and Strategic Priority Research Program of the Chinese Academy of Science (NO.Y929731103) and State Key Laboratory of Integrated Management of Pest Insects and Rodents, IOZ, CAS (NO.Y652751E03).

## AUTHOR CONTRIBUTIONS

Tao Wang, Resources, Data curation, Writing-original draft; Fengming Wu, Bowen Deng, Methodology; Kai Shi, Biyang Jing, Software; Baoxu Ma, Jing Li, Data curation; Chuan Zhou, Conceptualization, Funding acquisition, Writing-review and editing.

---

**Figure 1-figure supplement 1.**
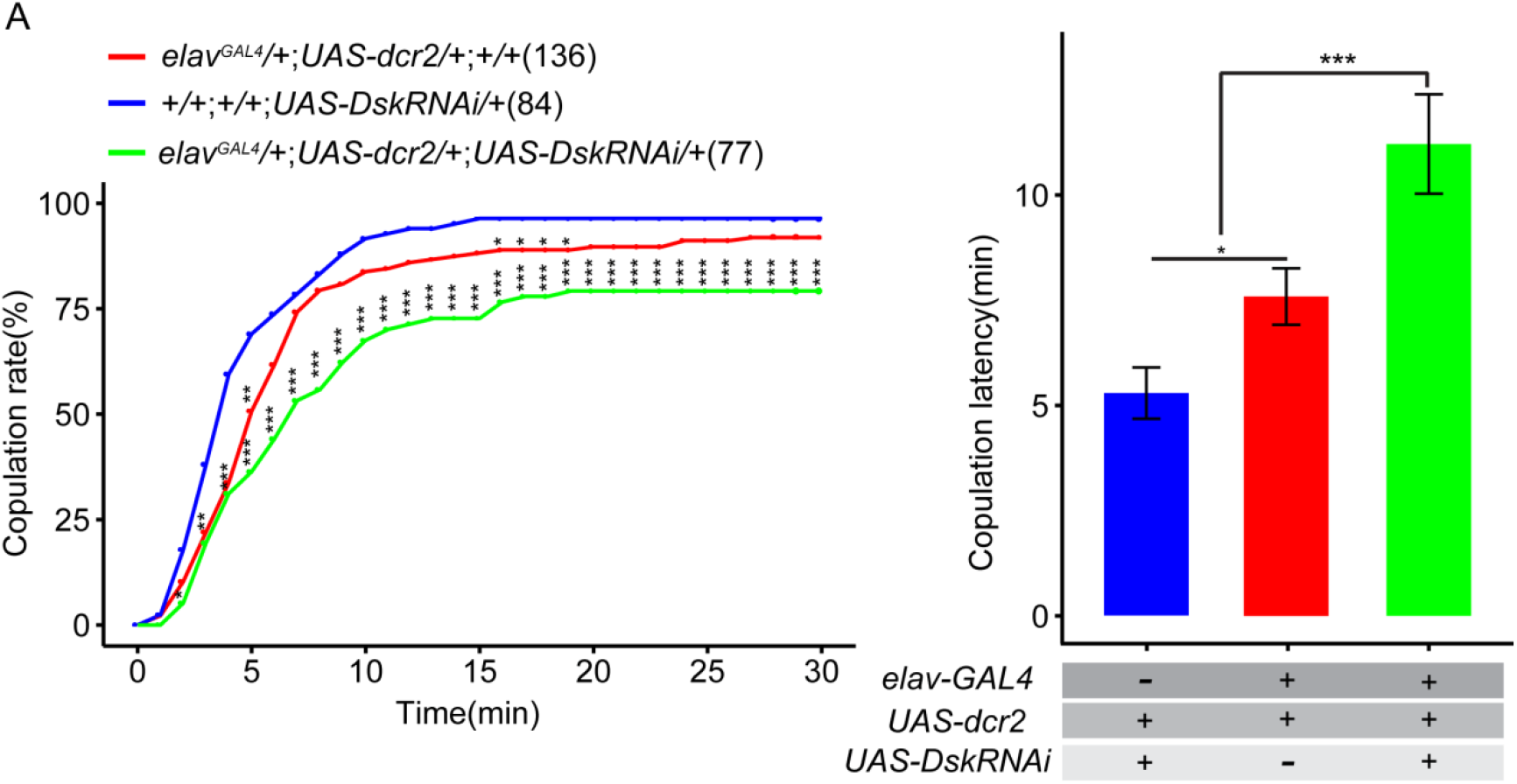
Knockdown of Dsk reduce female receptivity. (A) Knockdown of *Dsk* significantly decreased copulation rate and prolonged copulation latency compared with controls. *UAS-DskRNAi* was driven by *elav^GAL4^;UAS-dcr2*. The number of female flies paired with wild-type males analyzed is displayed in parentheses. And there are same numbers in two parameters. NS indicates no significant difference (chi-square test).

**Figure 1-figure supplement 2.**
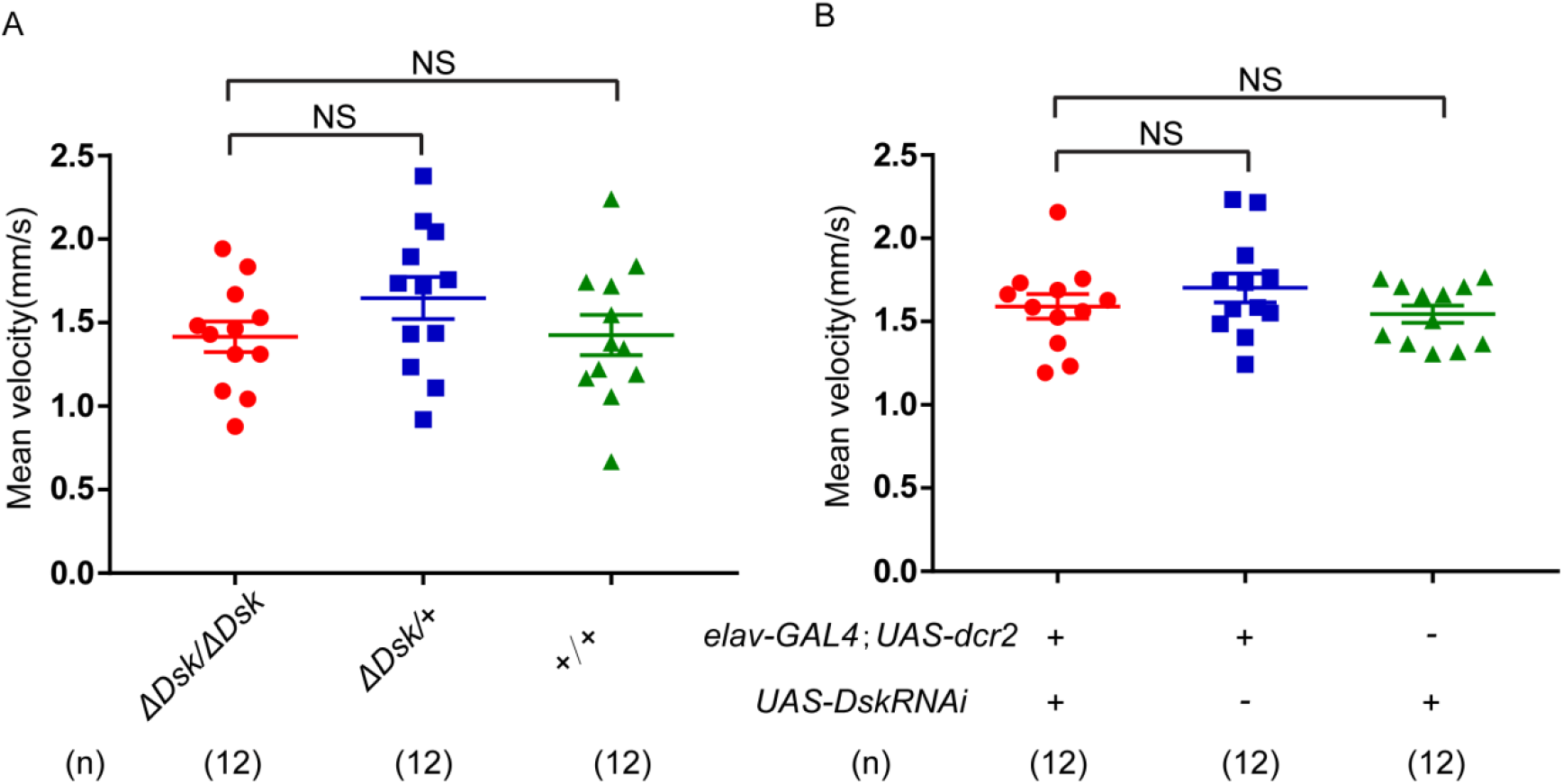
Locomotion behavior of Dsk mutant and Dsk RNAi in female. (A-B) Mean velocity had no significant change in *Dsk* mutant females (A) and *Dsk* RNAi females (B). Error bars indicate SEM. NS indicates no significant difference (Kruskal-Wallis and post-hoc Mann-Whitney U tests or post-hoc Student’s T-test).

**Figure 1-figure supplement 3.**
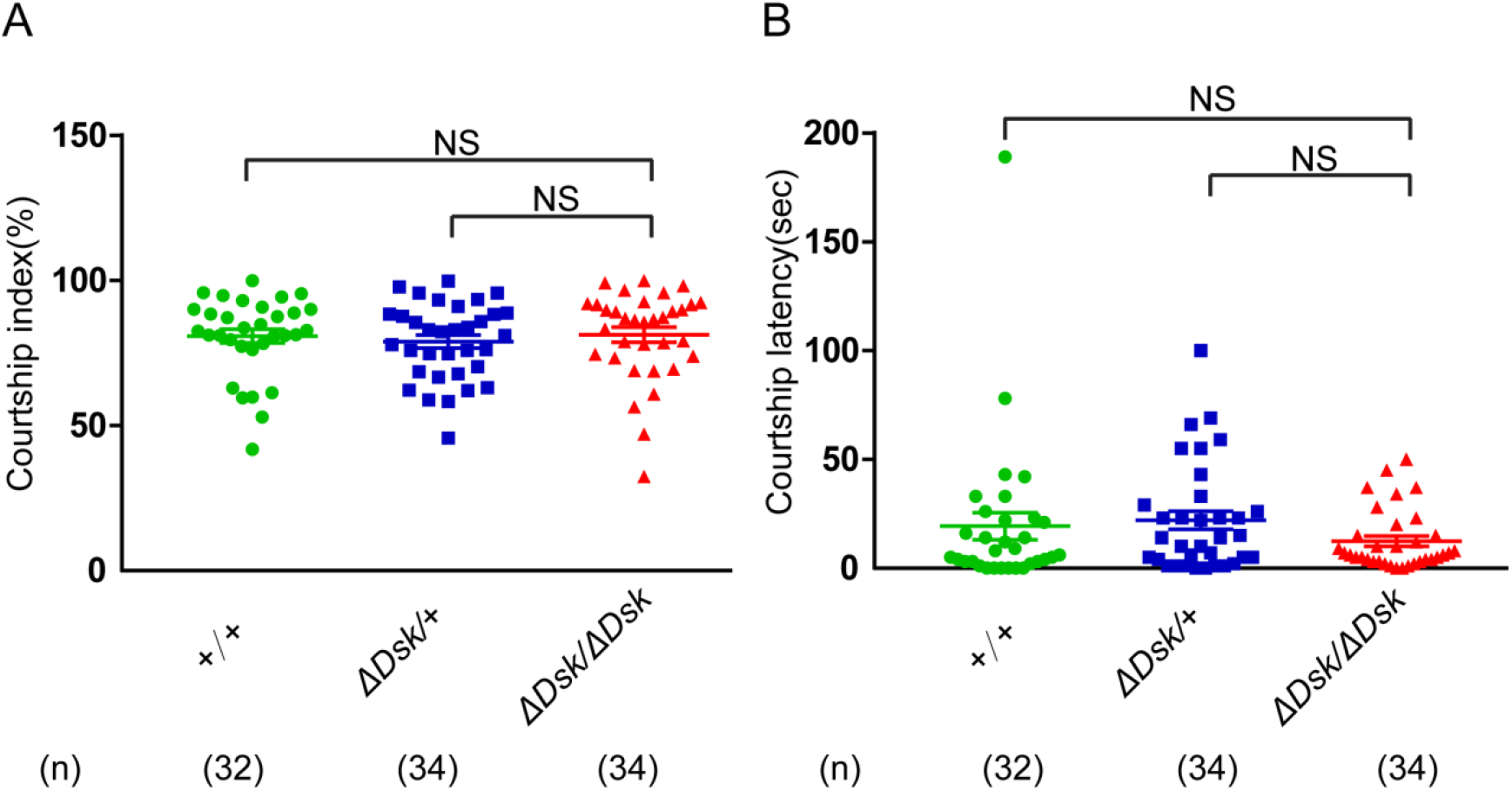
Courtship behavior of wild-type males paired with females of indicated genotypes. (A-B) Courtship index (%) and courtship latency (sec) in wild-type males paired with tested genotypes. Females for *ΔDsk/ΔDsk*, *+/ΔDsk*, *+/+* were used. Error bars indicate SEM. NS indicates no significant difference (Kruskal-Wallis and post-hoc Mann-Whitney U tests or post-hoc Student’s T-test).

**Figure 1-figure supplement 4.**
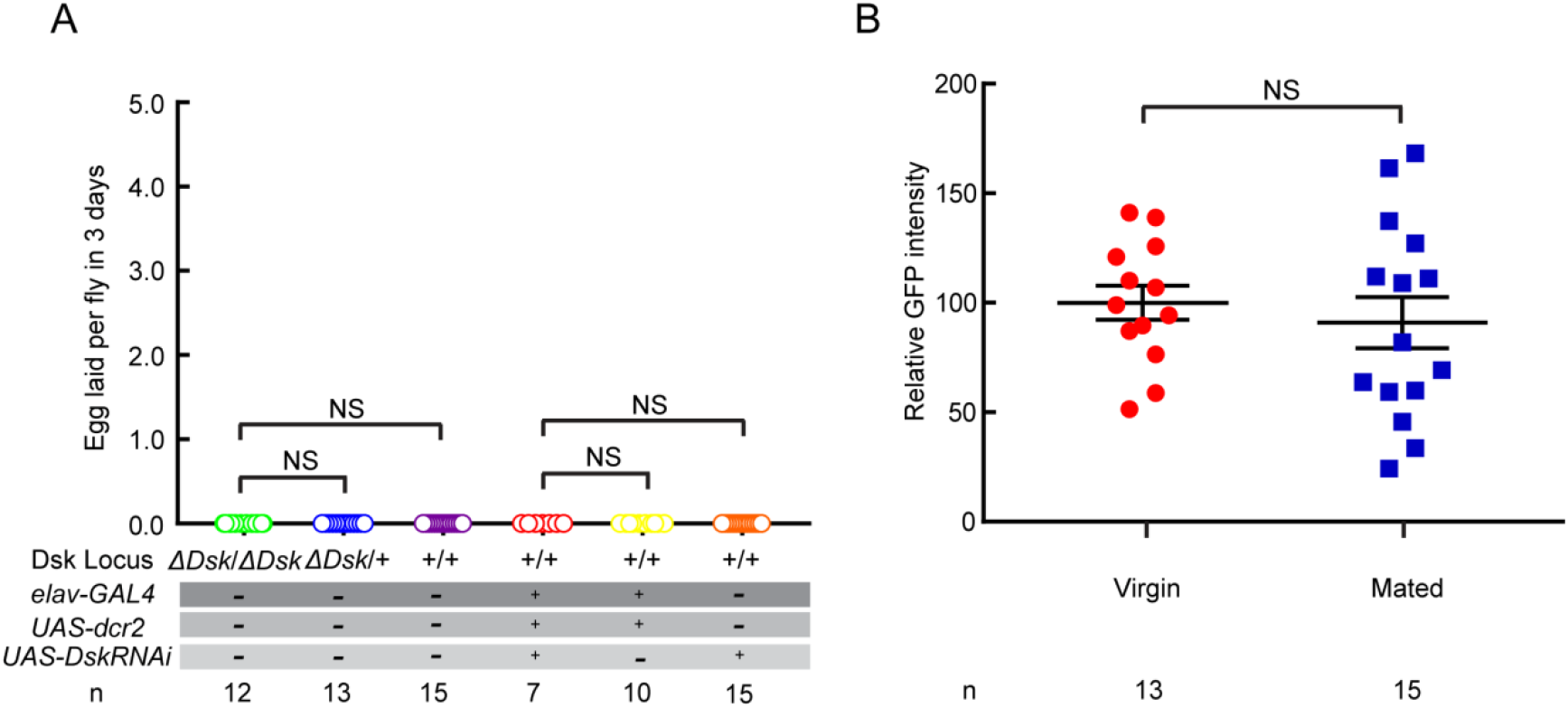
Dsk is not required for egg laying. (A) The number of eggs laid by virgin females within 3 days after mutation or RNAi of *Dsk*. (B) Ca^+^ activity of DSK neurons in virgin (red) and mated female (blue) brains detected by TRIC approach. Genotype: *UAS-IVS-mCD8∷RFP,LexAop2-mCD8∷GFP/+; nSyb-MKII∷nlsLexADBDo/+;Dsk^GAL4^/UAS-p65AD∷CaM.* NS indicates no significant difference (Kruskal-Wallis and post-hoc Mann-Whitney U tests).

**Figure 1-figure supplement 5.**
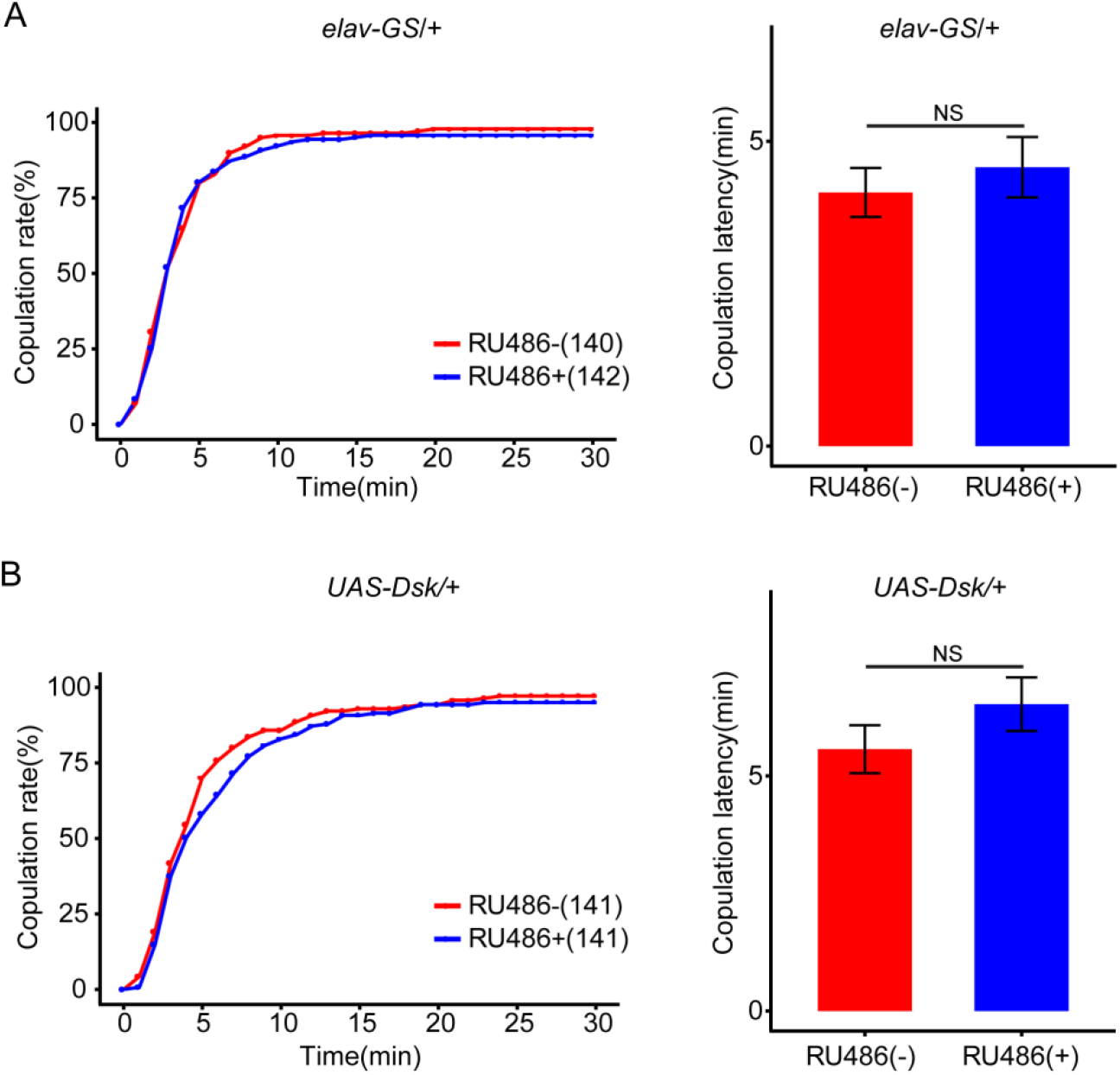
Control for overexprssion of Dsk. (A-B) The controls with either *elav-GS* alone or *UAS-Dsk* alone did not alter the copulation rate and copulation latency at feeding RU486 relative to without feeding RU486. The number of female flies paired with wild-type males analyzed is displayed in parentheses. And there are same numbers in two parameters. NS indicates no significant difference (chi-square test).

**Figure 1-figure supplement 6.**
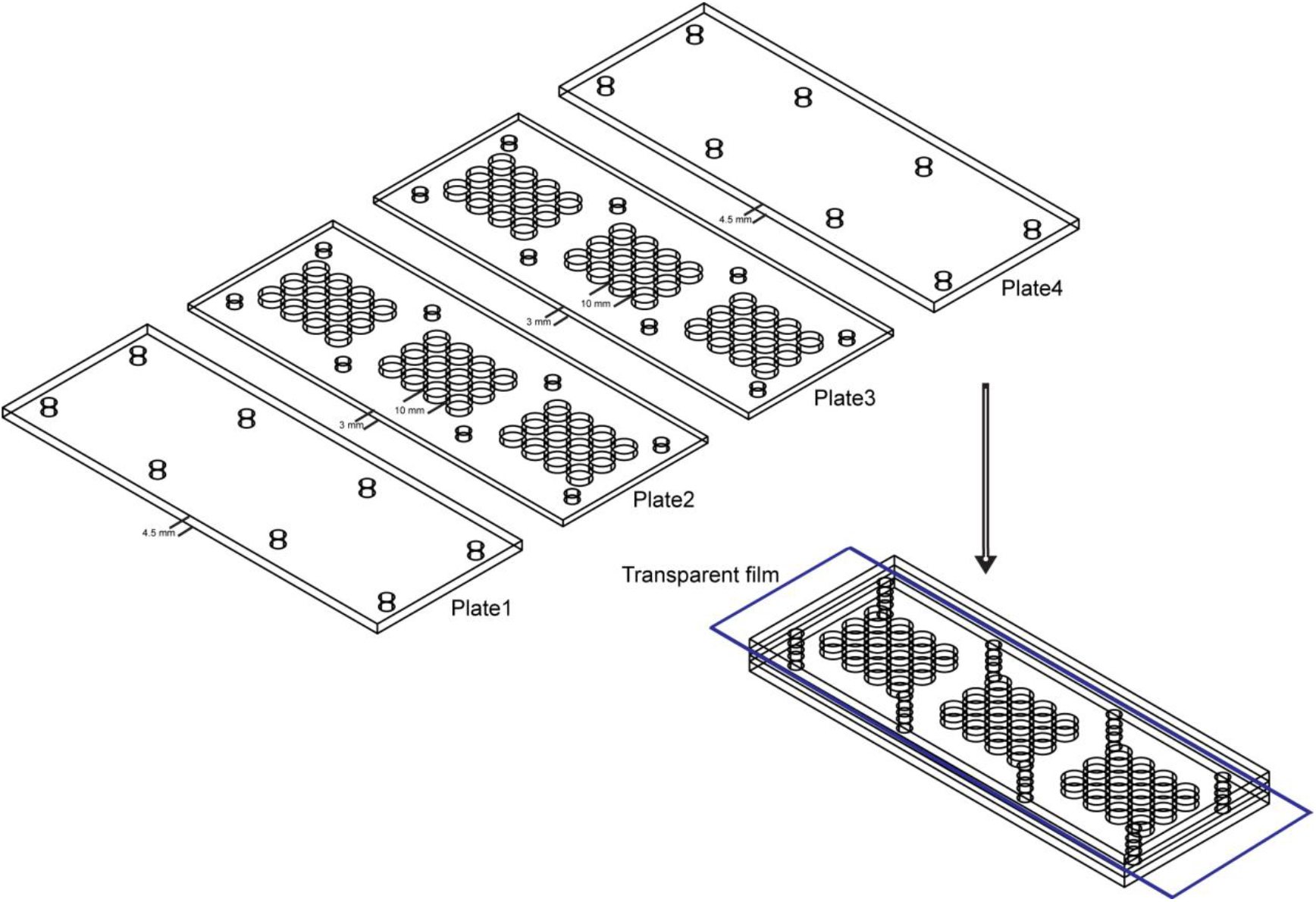
Behavior arena used in mating behavior assay. The mating arena contains four acrylic plates. The top (Plate1) and bottom (Plate4) are made up of acrylic plates of a thickness of 4.5mm. And the middle two layers (Plate2 and Plate3) are made up of acrylic plates with 36 cylindrical arenas (diameter: 10mm; height of each plates: 3mm). A removable transparent film was placed between Plate2 and Plate3 to separate the two flies and the film was removed to allow the pair of a test female and a wild-type male to contact.

**Figure 2-figure supplement 1.**
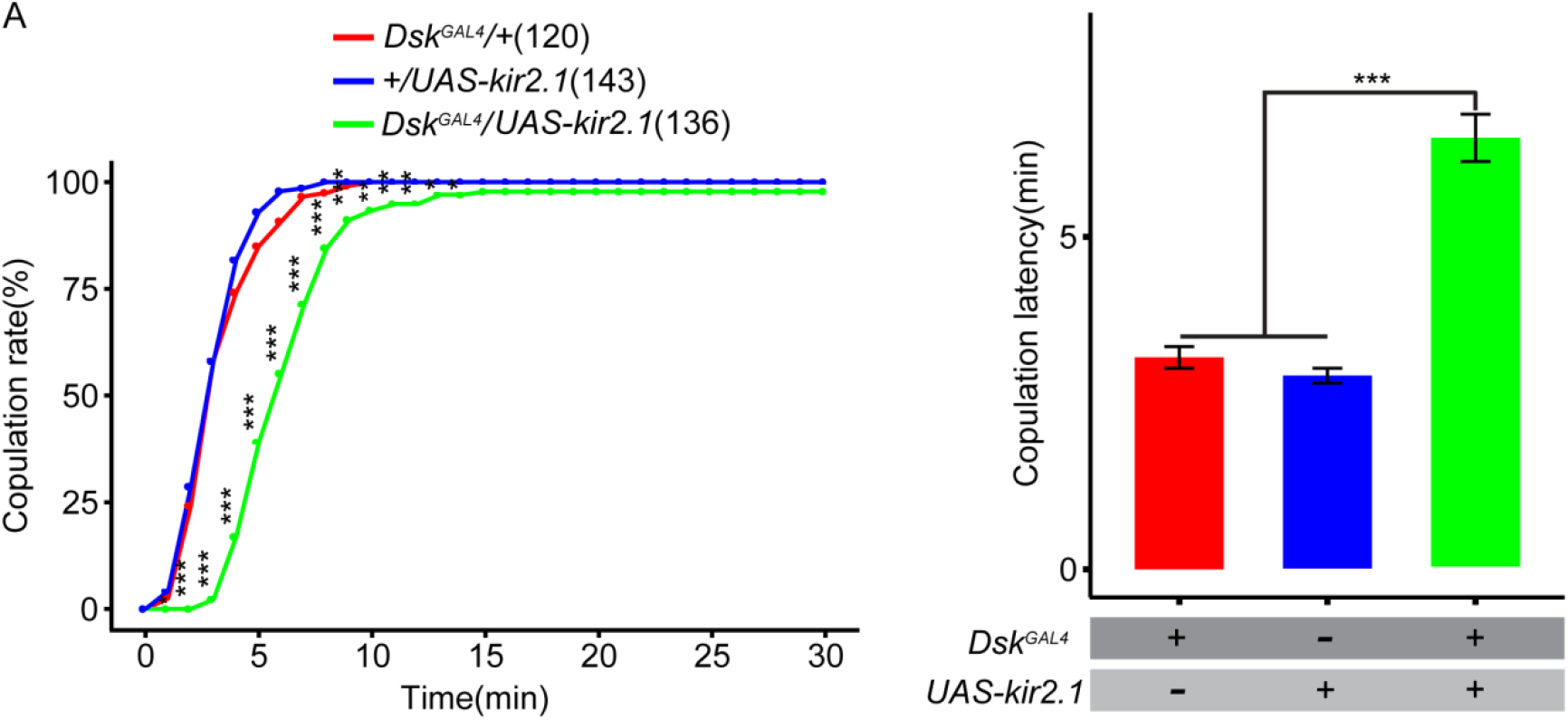
Effect of inactivation of DSK neurons on female receptivity. (A) Inactivation of DSK neurons significantly decreased copulation rate and prolonged copulation latency compared with controls. *Dsk^GAL^*^4^ driving *UAS-kir2.1* inactivates DSK neurons. The number of female flies paired with wild-type males analyzed is displayed in parentheses. And there are same numbers in two parameters. *p<0.05, **p<0.01, ***p<0.001, NS indicates no significant difference (chi-square test).

**Figure 2-figure supplement 2.**
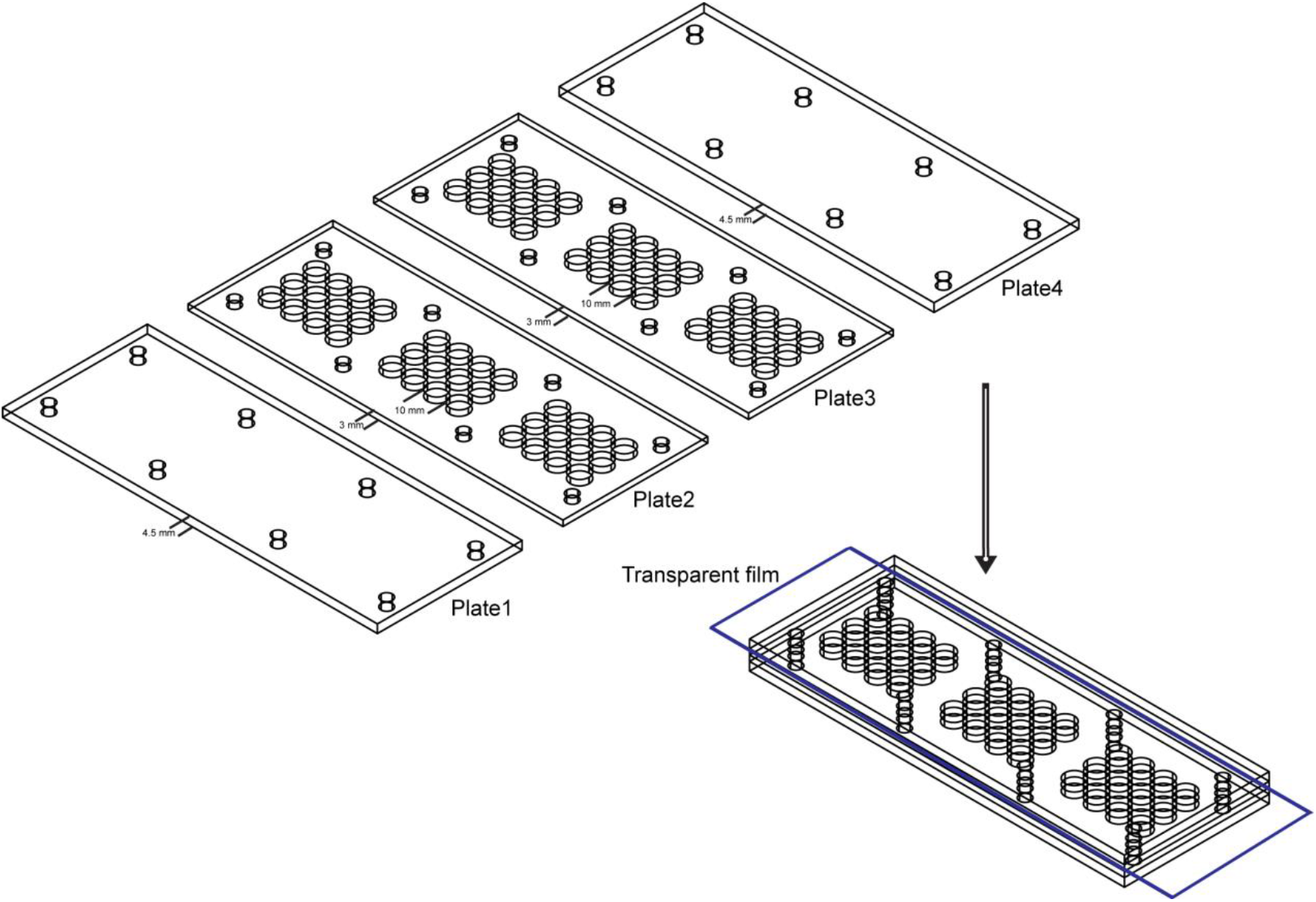
Behavior arena used in mating behavior assay. The mating arena contains four acrylic plates. The top (Plate1) and bottom (Plate4) are made up of acrylic plates of a thickness of 4.5mm. And the middle two layers (Plate2 and Plate3) are made up of acrylic plates with 36 cylindrical arenas (diameter: 10mm; height of each plates: 3mm). A removable transparent film was placed between Plate2 and Plate3 to separate the two flies and the film was removed to allow the pair of a test female and a wild-type male to contact.

**Figure 3-supplement 1.**
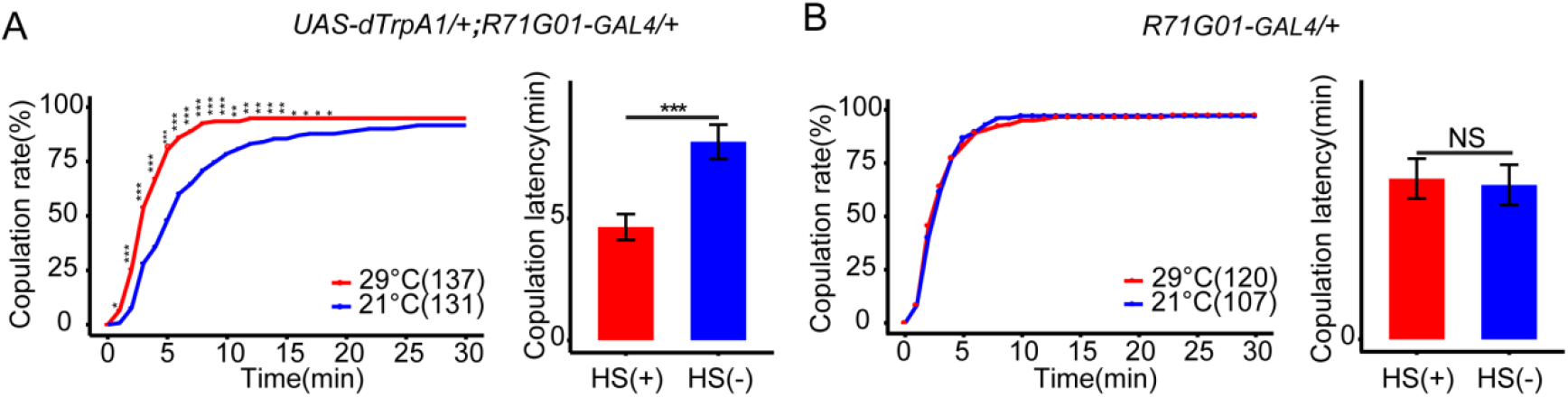
Effect of R71G01GAL4 neurons on female receptivity. (A) Activation of *R71G01GAL4* neurons significantly increased copulation rate and shortened copulation latency at 29°C relative to 21°C. *R71G01GAL*4 driving *UAS-dTrpA1* activated *R71G01GAL4* neurons at 29°C. (B) The controls with *R71G01GAL*4 alone did not alter the copulation rate and copulation latency at 29°C relative to 21°C. Activation of neuron is represented by (HS+) and control is represented by (HS-). The number of female flies paired with wild-type males analyzed is displayed in parentheses. And there are same numbers in two parameters. *p<0.05, **p<0.01, ***p<0.001, NS indicates no significant difference (chi-square test).

**Figure 3-supplement 2.**
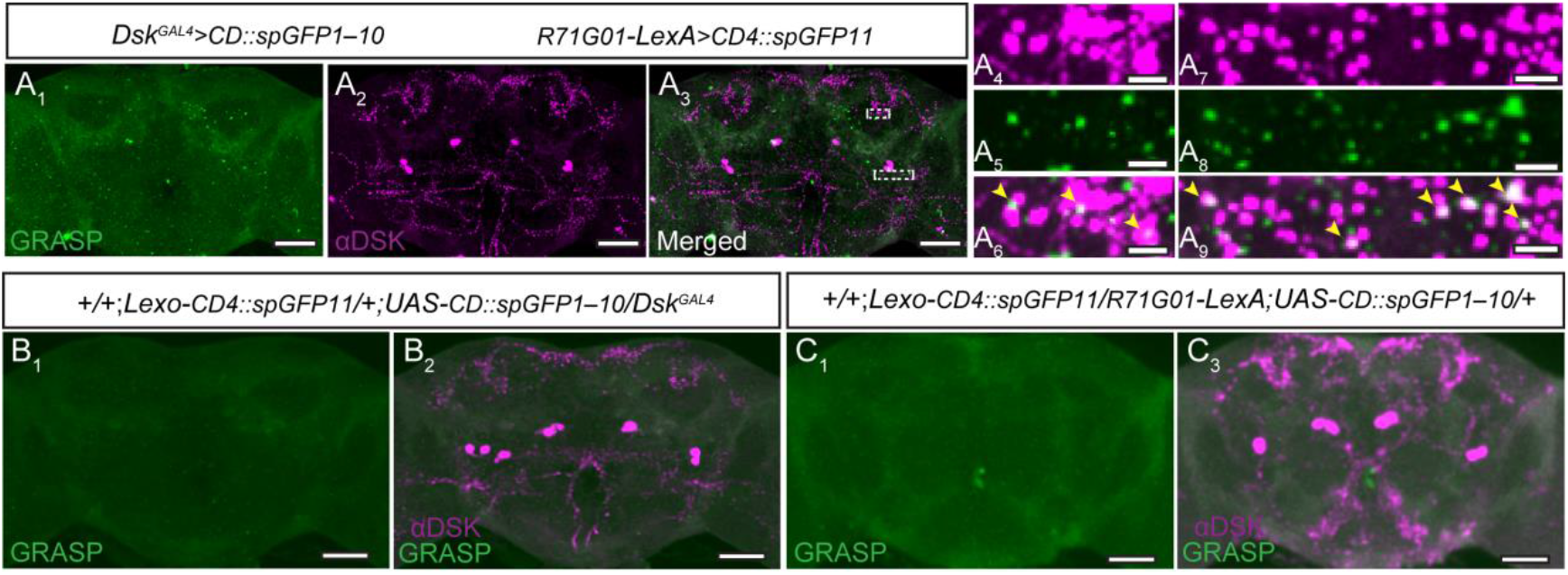
Potential connection between R70G01GAL4 neurons and DSK neurons was detected by GRASP method. (A) The recombinant GFP signals (GRASP) (A_1_) revealed synaptic contacts between *R70G01GAL4* neurons and DSK neurons. *Dsk^GAL^*^4^ expresses *CD4∷spGFP11*, R*71G01-LexA* expresses *CD4∷spGFP1-10*. (A_2_) anti-DSK antibody staining. (A_3_) Merge. Magnification of white boxed region in (A_3_) is shown in (A_4_-A_6_) and (A_7_-A_9_). Yellow arrowheads indicated GRASP signal co-localized with synaptic boutons revealed by anti-DSK antibody staining. (B-C) No recombinant GFP signals (GRASP) were observed in flies with either *Dsk^GAL^*^4^ alone (B_1_) or R*71G01-LexA* alone (C_1_). Scale bars are 50μm in (A-C), 5μm in (A_3_-A_9_).

**Figure 3-supplement 3.**
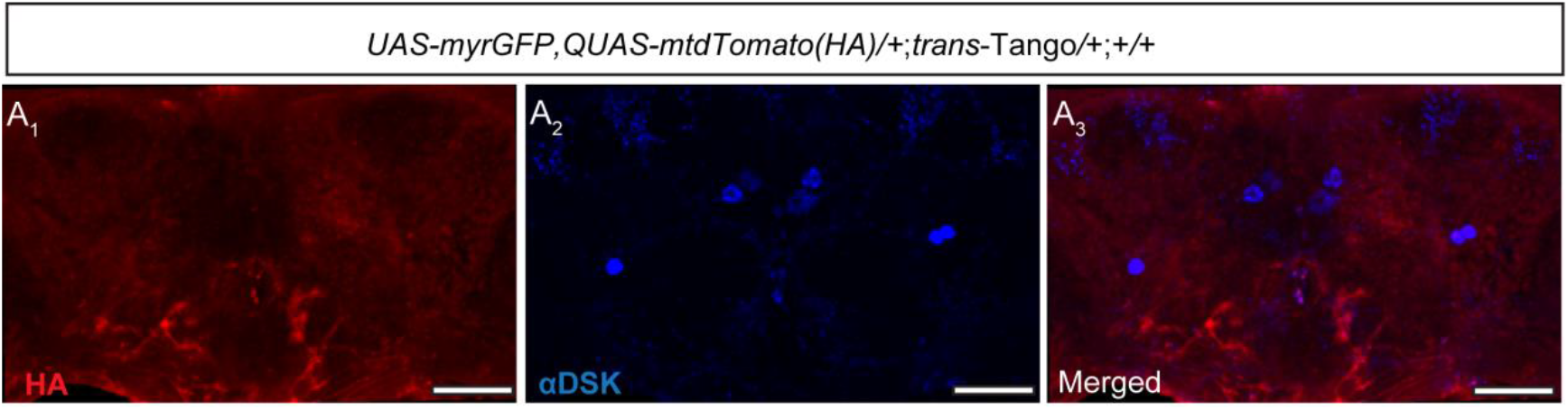
Control for the trans-Tango experiment. (A) No postsynaptic signals (anti-HA, red) (A_1_) were detected in the central brain without a Gal4 driver. Cell bodies of Dsk were stained with anti-DSK (blue) (A_2_). Merge (A_3_) Scale bars: 50μm.

**Figure 3-figure supplement 4.**
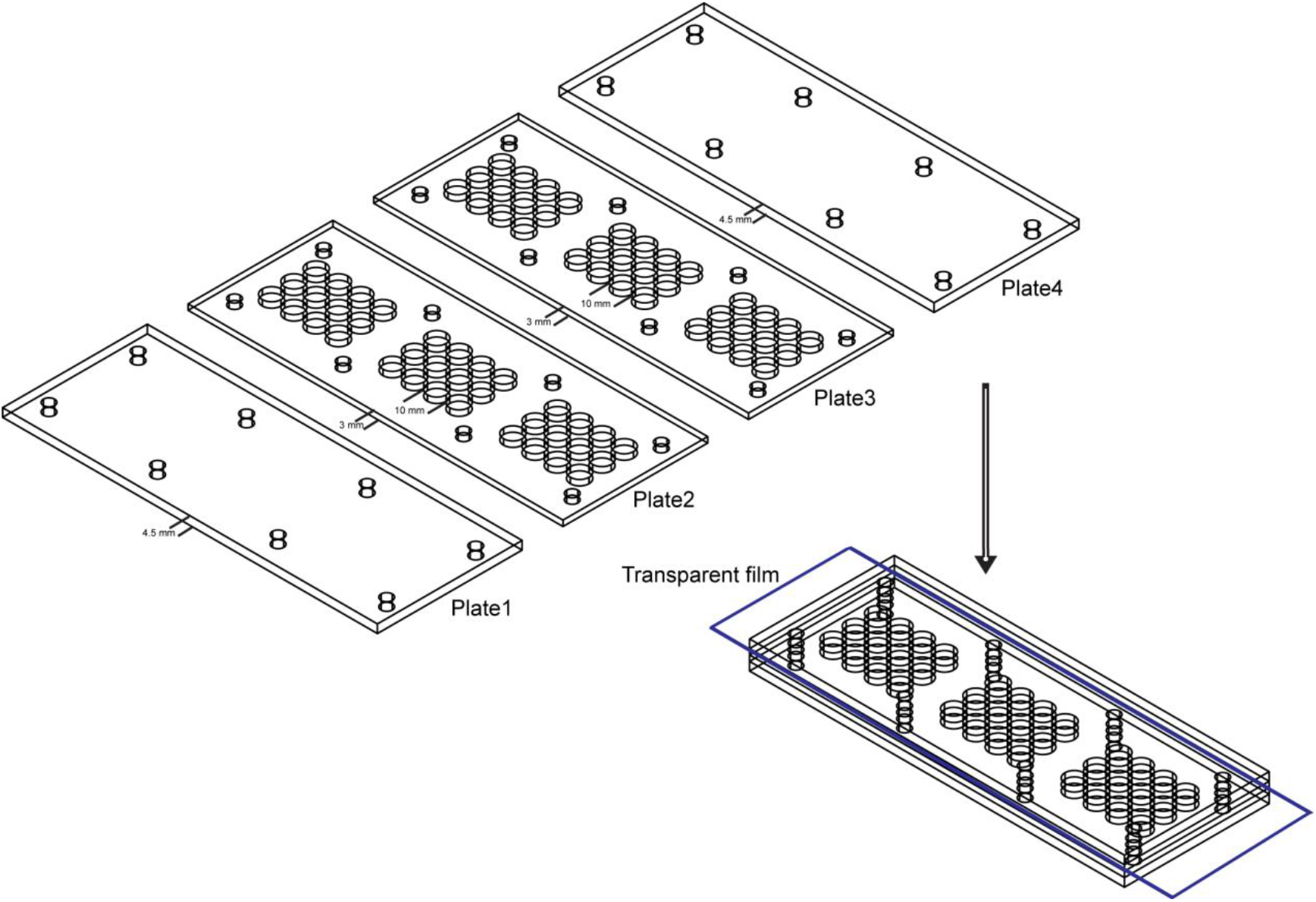
Behavior arena used in mating behavior assay. The mating arena contains four acrylic plates. The top (Plate1) and bottom (Plate4) are made up of acrylic plates of a thickness of 4.5mm. And the middle two layers (Plate2 and Plate3) are made up of acrylic plates with 36 cylindrical arenas (diameter: 10mm; height of each plates: 3mm). A removable transparent film was placed between Plate2 and Plate3 to separate the two flies and the film was removed to allow the pair of a test female and a wild-type male to contact.

**Figure 5-figure supplement 1.**
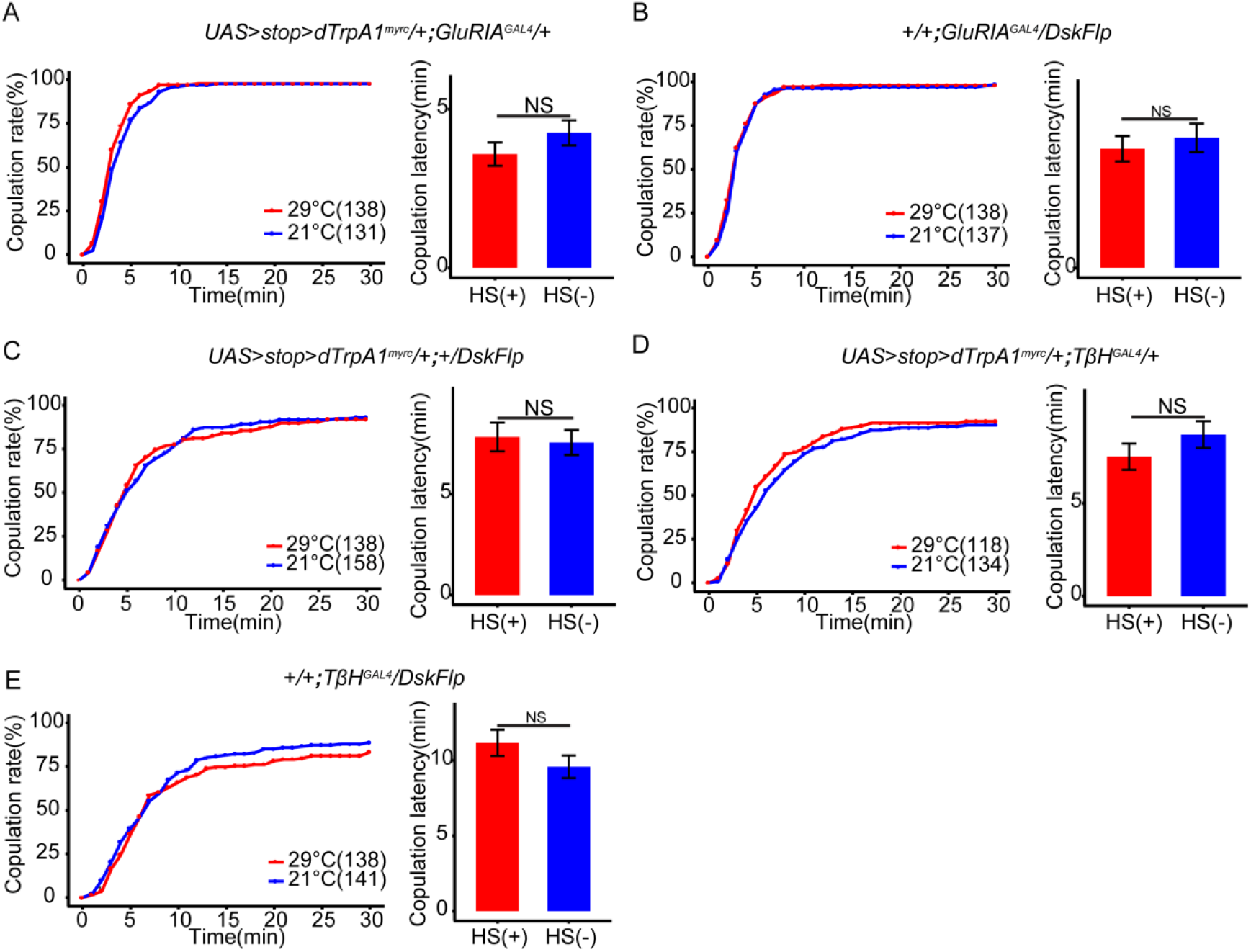
Control for activation of intersectional expression neurons (DSK-M and DSK-L). (A-C) The controls of activation of DSK-M neurons did not alter the copulation rate and shortened copulation latency at 29°C relative to 21°C in the absence of *DskFlp* (A), *UAS>stop>dTrpA^myrc^* (B) and *GluRIA^GAL4^* (C). (D-E) The controls of activation of DSK-L neurons also did not alter the copulation rate and shortened copulation latency at 29°C relative to 21°C in the absence of *TβH^GAL4^* (C), *DskFlp* (D), *UAS>stop>dTrpA^myrc^* (E). The number of female flies paired with wild-type males analyzed is displayed in parentheses. And there are same numbers in two parameters. NS indicates no significant difference (chi-square test).

**Figure 5-figure supplement 2.**
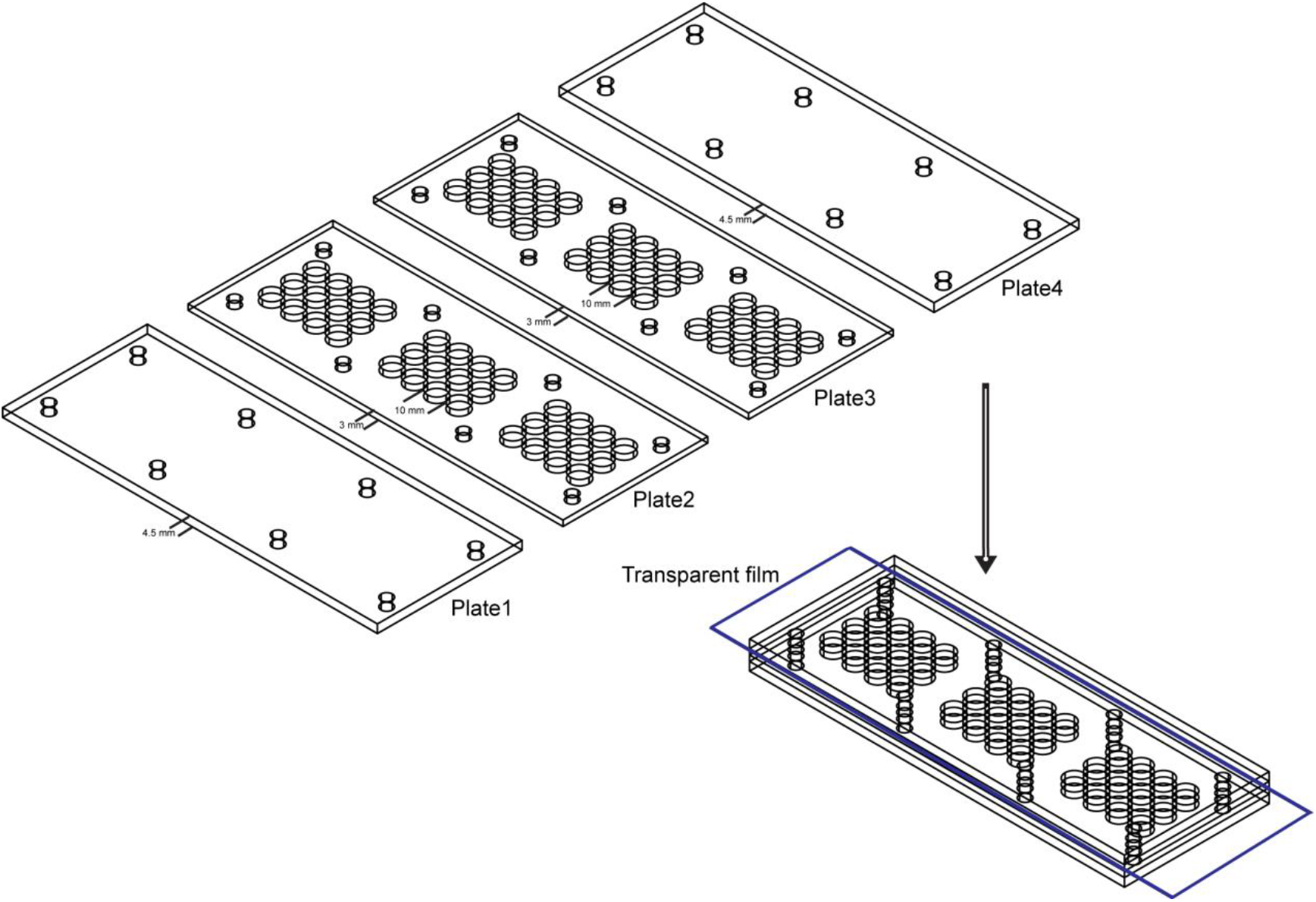
Behavior arena used in mating behavior assay. The mating arena contains four acrylic plates. The top (Plate1) and bottom (Plate4) are made up of acrylic plates of a thickness of 4.5mm. And the middle two layers (Plate2 and Plate3) are made up of acrylic plates with 36 cylindrical arenas (diameter: 10mm; height of each plates: 3mm). A removable transparent film was placed between Plate2 and Plate3 to separate the two flies and the film was removed to allow the pair of a test female and a wild-type male to contact.

**Figure 6-figure supplement 1.**
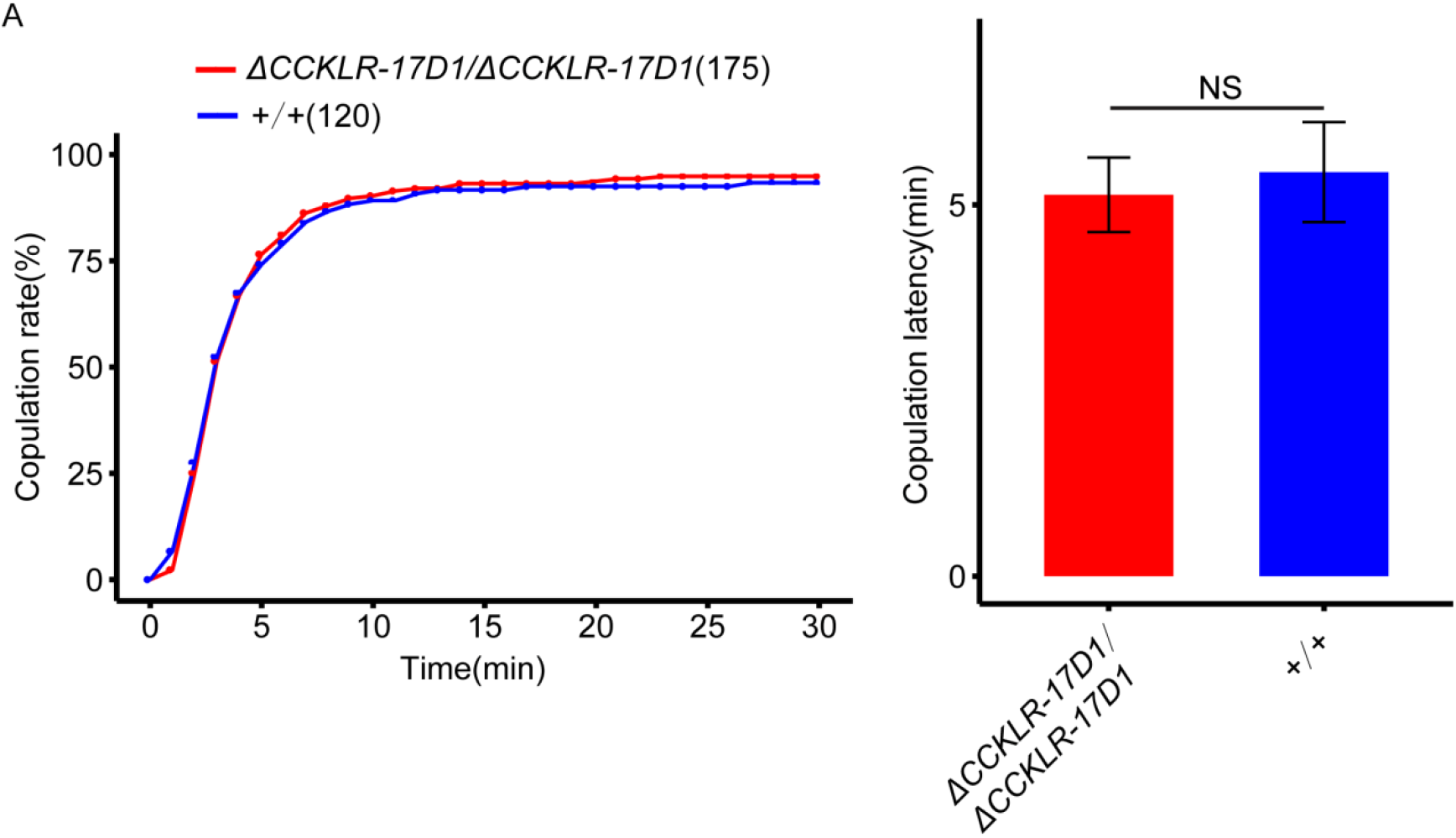
Effect of CCKLR-17D1 mutant on female receptivity. (A) *CCKLR-17D1* mutant did not alter the copulation rate and copulation latency compared with wild-type females. The number of female flies paired with wild-type males analyzed is displayed in parentheses. And there are same numbers in two parameters. NS indicates no significant difference (chi-square test).

**Figure 6-figure supplement 2.**
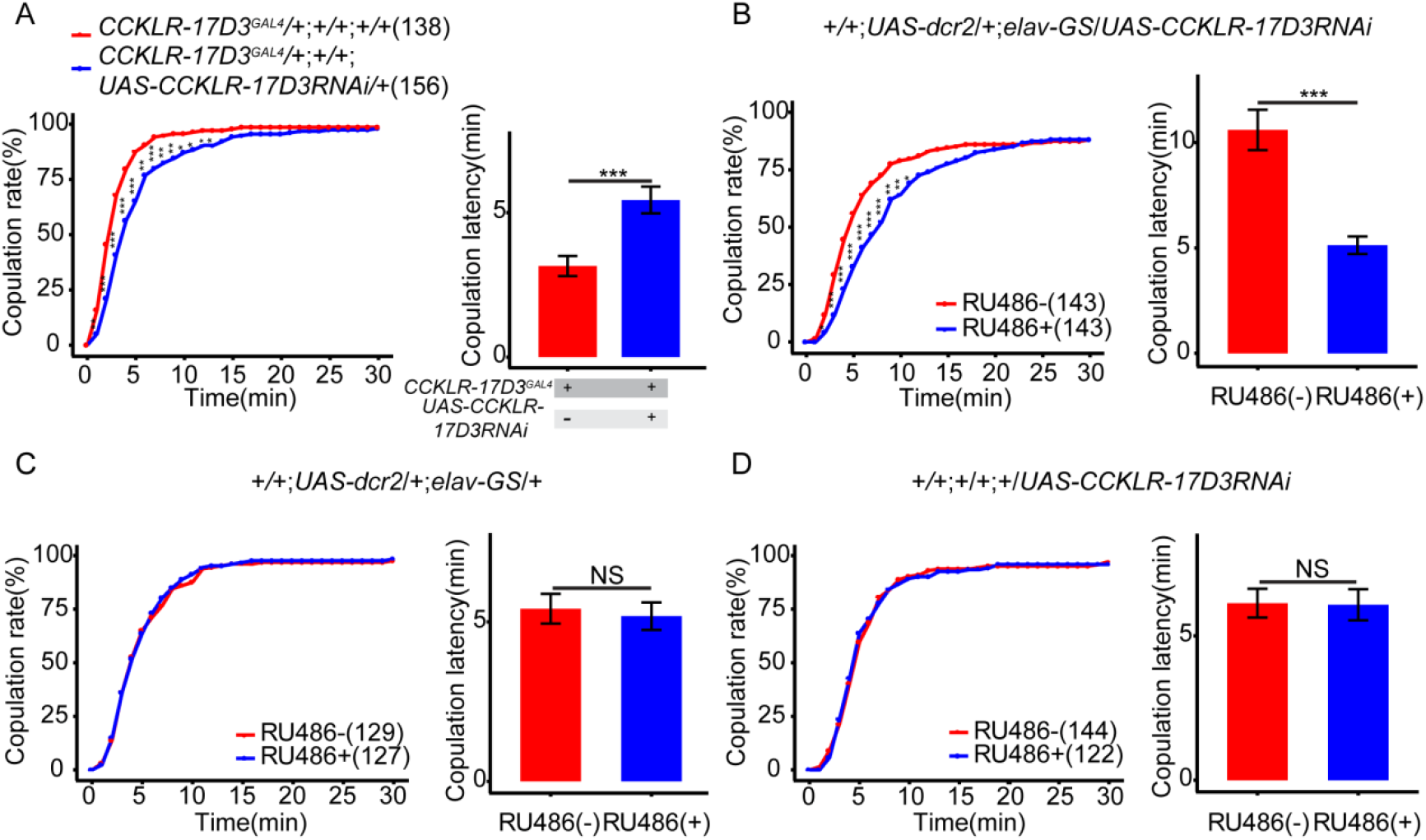
Effect of CCKLR-17D3 RNAi on female receptivity. (A) Knockdown of *CCKLR-17D3* significantly decreased copulation rate and prolonged copulation latency compared with controls. *UAS-CCKLR-17D3* RNAi was driven by *CCKLR-17D3^GAL4^*. (B) Conditional knockdown of *CCKLR-17D3* after feeding RU486 significantly decreased copulation rate and prolonged copulation latency compared without feeding RU486. *UAS-CCKLR-17D3* RNAi was driven by *UAS-dcr2;elav-GS*. (C-D) The controls with either *UAS-dcr2;elav-GS* alone or *UAS-CCKLR-17D3* RNAi alone did not alter the copulation rate and copulation latency at feeding RU486 relative to without feeding RU486. The number of female flies paired with wild-type males analyzed is displayed in parentheses. And there are same numbers in two parameters. *p<0.05, **p<0.01, ***p<0.001, NS indicates no significant difference (chi-square test).

**Figure 6-figure supplement 3.**
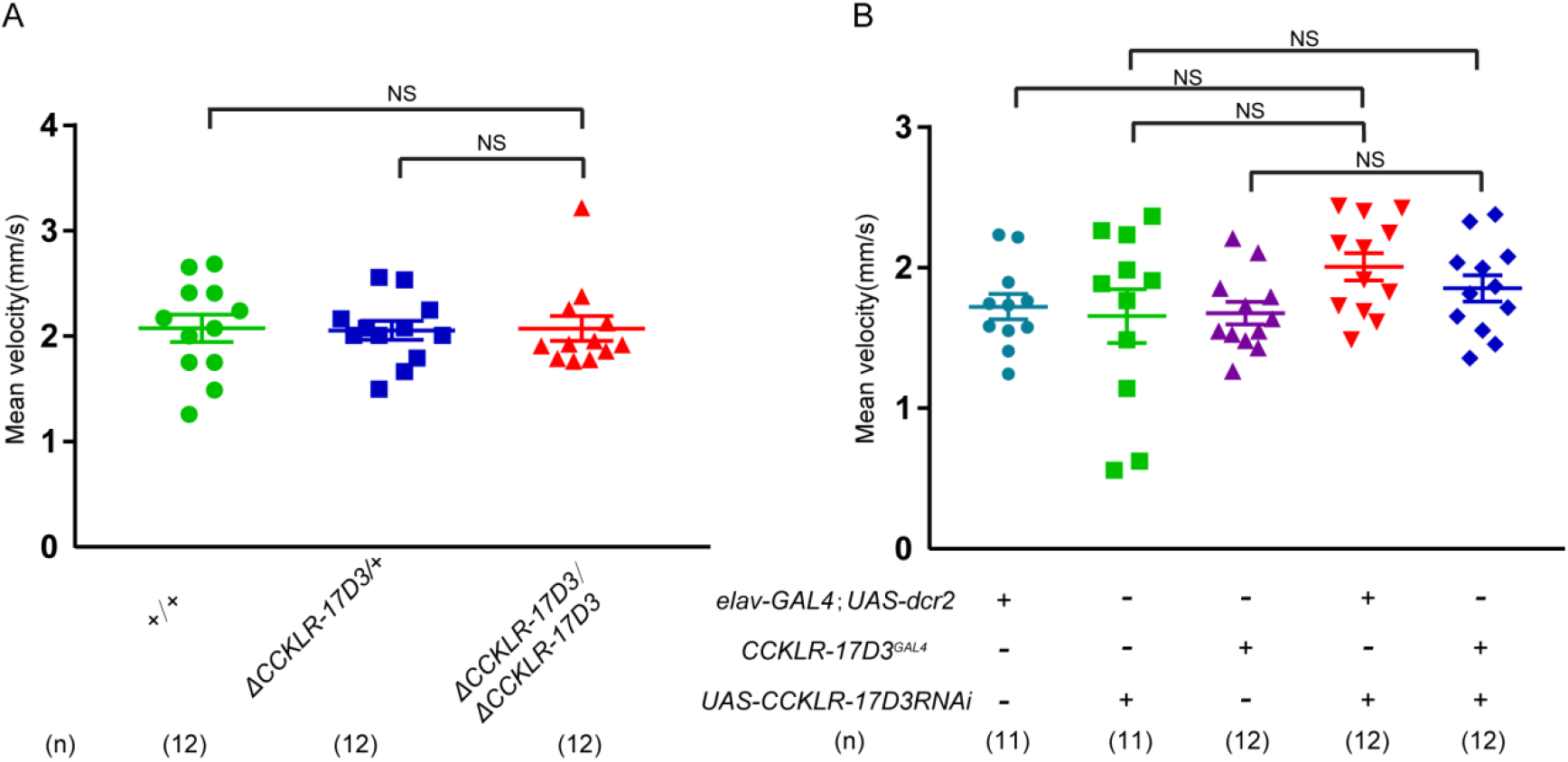
Locomotion behavior of Δ*CCKLR-17D3 mutant and CCKLR-17D3* RNAi in female. (A-B) Mean velocity had no significant changes in *CCKLR-17D3* mutant females (A) and *CCKLR-17D3* RNAi females (B). Error bars indicate SEM. NS indicates no significant difference (Kruskal-Wallis and post-hoc Mann-Whitney U tests or post-hoc Student’s T-test).

**Figure 6-figure supplement 4.**
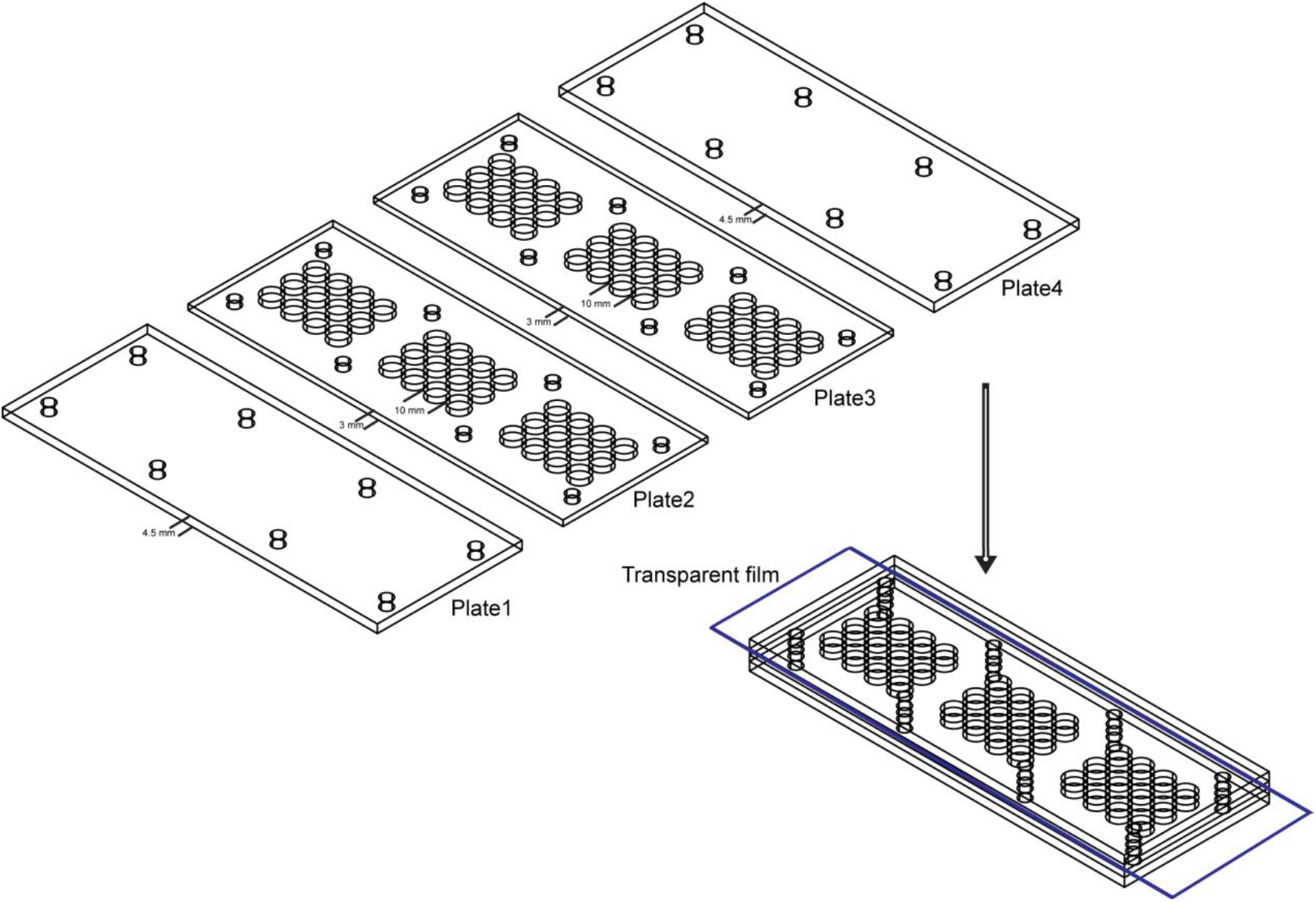
Behavior arena used in mating behavior assay. The mating arena contains four acrylic plates. The top (Plate1) and bottom (Plate4) are made up of acrylic plates of a thickness of 4.5mm. And the middle two layers (Plate2 and Plate3) are made up of acrylic plates with 36 cylindrical arenas (diameter: 10mm; height of each plates: 3mm). A removable transparent film was placed between Plate2 and Plate3 to separate the two flies and the film was removed to allow the pair of a test female and a wild-type male to contact.

